# *CDK4* is co-amplified with either *TP53* promoter gene fusions or *MDM2* through distinct mechanisms in osteosarcoma

**DOI:** 10.1101/2024.03.13.584810

**Authors:** Karim H Saba, Valeria Difilippo, Emelie Styring, Jenny Nilsson, Linda Magnusson, Hilda van den Bos, Diana C. J. Spierings, Floris Foijer, Michaela Nathrath, Felix Haglund de Flon, Daniel Baumhoer, Karolin H Nord

## Abstract

Amplification of the *MDM2* and *CDK4* genes on chromosome 12 is commonly associated with low-grade osteosarcomas. In this study, we conducted high-resolution genomic and transcriptomic analyses on 33 samples from 25 osteosarcomas, encompassing both high- and low-grade cases with *MDM2* and/or *CDK4* amplification. We identified four major subgroups: (i) low-grade osteosarcoma with chromosome 12 amplicons as the sole acquired alteration, (ii) high- and low-grade tumours with *CDK4* and *MDM2* amplification along with few changes affecting other chromosomes, (iii) high-grade osteosarcomas with heavily rearranged genomes including either *CDK4* and *MDM2* amplification or (iv) *CDK4* amplification and *TP53* structural alterations. The amplicons involving *MDM2* exhibited signs of an initial chromothripsis event affecting chromosome 12. In contrast, there was no indication of a chromothripsis event on chromosome 12 in *TP53*-rearranged cases. Instead, the initial disruption of the *TP53* locus resulted in breakage and repair processes that co-amplified the *CDK4* locus. Furthermore, our investigation revealed recurring promoter swapping events that involved the regulatory regions of the *FRS2*, *PLEKHA5*, and *TP53* genes. These events led to the ectopic expression of partner genes, with the *ELF1* gene being upregulated by the *FRS2* and *TP53* promoter regions, respectively, in two distinct cases.

## Introduction

Osteosarcoma exhibits distinctive subgroups based on clinical, morphological and genetic characteristics. The most prevalent subtype is conventional osteosarcoma, a high-grade tumour necessitating multimodal therapy. Conventional osteosarcoma typically affects children and adolescents but may manifest at any age. The overall survival is up to 70% for patients with localised disease and good response to neoadjuvant chemotherapy, but the prognosis markedly deteriorates for patients with metastatic or recurrent tumours [1–3]. In contrast, low-grade osteosarcoma is rare and generally has a favourable prognosis, with a five-year survival rate of 90% [4, 5]. However, there is a potential for dedifferentiation, leading to the transformation into a high-grade tumour with a prognosis similar to that of conventional osteosarcoma [6, 7]. Predominantly affecting adults, with a slight female predominance, low-grade osteosarcoma is further categorised based on tumour site in the bone; low-grade central osteosarcoma originates within the medullary cavity, and parosteal osteosarcoma emerges on the cortical surface [4, 5].

A genetic hallmark of parosteal osteosarcoma is the presence of supernumerary ring or marker chromosomes, often observed as the sole cytogenetically visible alteration [8–10]. These structures encompass amplified material from chromosomal region 12q13-15, housing genes such as *CDK4* and *MDM2,* and occasionally featuring gained or amplified material from chromosome arm 12p as well [8, 9, 11]. Although studies on low-grade central osteosarcoma are limited, available information suggests the presence of similar amplicons in this subtype, albeit at a lower frequency [12, 13]. In contrast, conventional osteosarcoma typically exhibits genome-wide structural and numerical chromosome changes, accompanied by the loss of *TP53* function [14–18]. In approximately 10% of cases, these complex alterations involve the amplification of chromosome bands 12q13-15 [11].

In this study, we conducted comprehensive genomic and transcriptomic analyses on both high- and low-grade osteosarcomas exhibiting *MDM2* and/or *CDK4* amplification and verified our findings using longread whole-genome sequencing. Our aim was to explore whether additional genes within the chromosome 12 amplicons might play a role in tumour progression. Our findings unveiled a correlation between genome-wide alterations and the amplification pattern on chromosome 12. This led to the identification of distinct genetic subgroups, ranging from predominantly intact genomes featuring chromosome 12 amplicons as the sole acquired alteration, to increasingly complex genomes with both *CDK4* and *MDM2* amplification, or *CDK4* co-amplified with *TP53* structural variants. In addition to gene amplification, we observed promoter swapping events that impacted the expression of various known or putative oncogenes, with the transcription factor *ELF1* being recurrently affected.

## Results

We selected osteosarcomas based on subtype or *CDK4* and *MDM2* status, drawing cases from two distinct cohorts. Cohort 1 (*n* = 17) comprised a diverse group of osteosarcomas diagnosed at Skåne University Hospital or Karolinska University Hospital in Sweden over the last four decades. Our dataset included DNA copy number and/or RNA sequencing data, revealing amplification and/or relatively high gene expression levels of *CDK4* and/or *MDM2*. Additionally, four cases lacking such information were included due to their diagnosis being parosteal osteosarcoma. Cohort 2 consisted of 108 unselected conventional osteosarcomas for which DNA copy number analyses were available [18]. Among the 108 osteosarcomas, 5% exhibited amplification of *CDK4*, but not *MDM2*, and 3% displayed amplification of both *CDK4* and *MDM2*. In total, we included 33 samples from 25 osteosarcomas in the present study (Supplementary Table S1). The selected cases underwent further exploration using SNP array analysis, bulk tumour whole-genome mate pair and longread sequencing, whole-transcriptome sequencing, and single-cell genome sequencing analyses. Integration of this information with pre-existing genetic and clinical data identified four genetic subgroups.

### Group A *CDK4* and *MDM2* amplified osteosarcomas with no alteration outside of chromosome 12

Group A comprised three low-grade osteosarcomas (one low-grade central and two parosteal) with amplification of *CDK4* and *MDM2,* along with regions on chromosome arm 12p (Figures 1 and 2, Supplementary Figure S1 and Supplementary Table S1). There were no deletions affecting chromosome 12 and no alteration was observed on other chromosomes in bulk DNA. Single-cell whole-genome analysis of one case revealed potential subclones (Supplementary Figure S1B). However, the absence of recurrent alterations outside of chromosome 12 suggests that these changes may be attributed to technical artifacts or non-clonal events. The distribution and read orientation of sequencing reads from chromosome 12 indicated a chromothripsis event (Figure 3, Supplementary Figure S1) [19]. In summary, it is likely that Group A amplicons originated from an initial chromothripsis event affecting an extra copy of chromosome 12, followed by subsequent rounds of selective amplification through breakage-fusion-bridge cycles [20–22].

**Figure 1.**
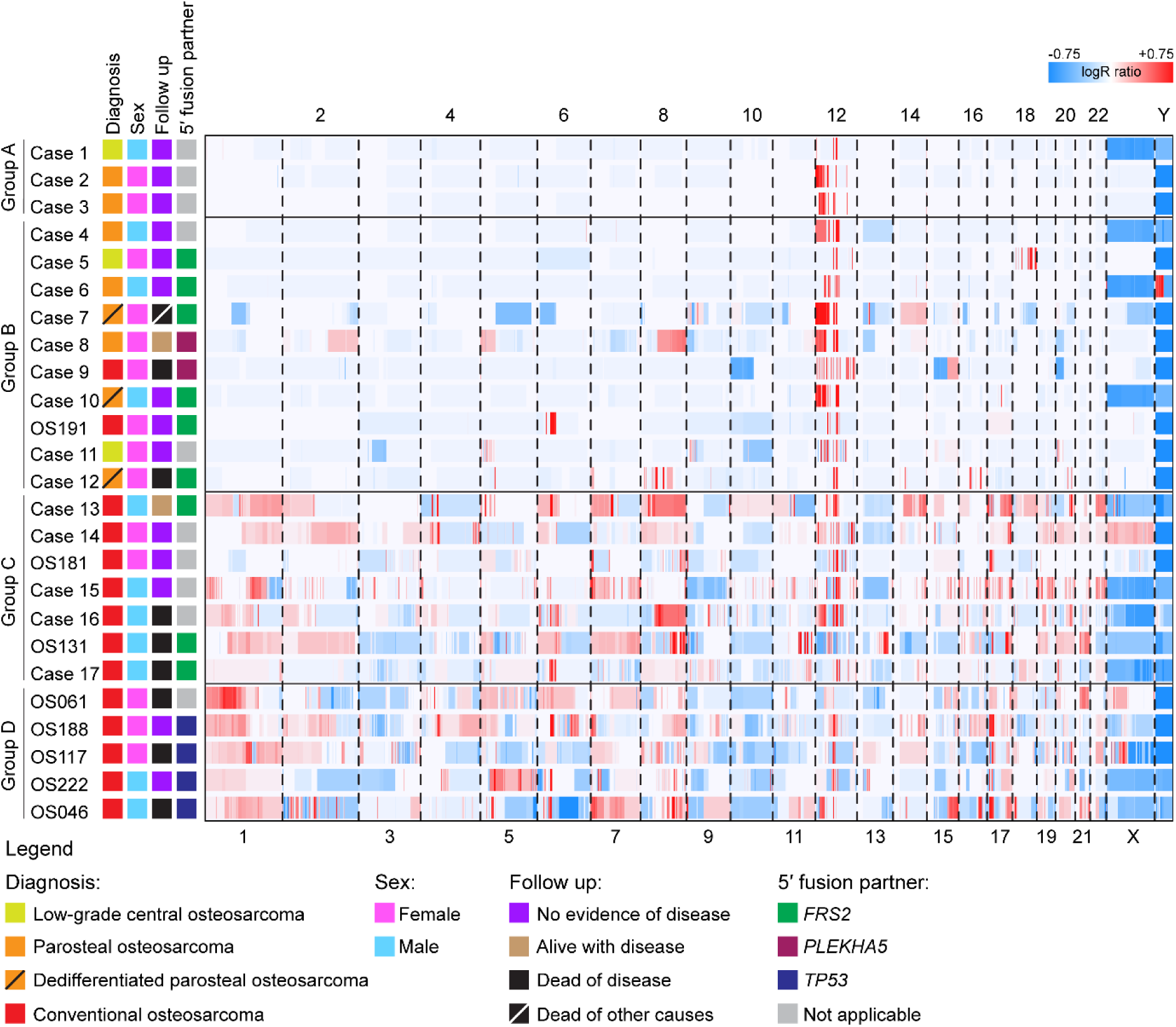
Whole-genome copy number heatmap of 25 osteosarcomas with amplification of 12q sequences. Genome-wide copy number gains (depicted in red) and losses (depicted in blue) were identified by copy number array analysis. The cases are categorised into four groups based on their genome-wide copy number alterations and *MDM2* status.

**Figure 2.**
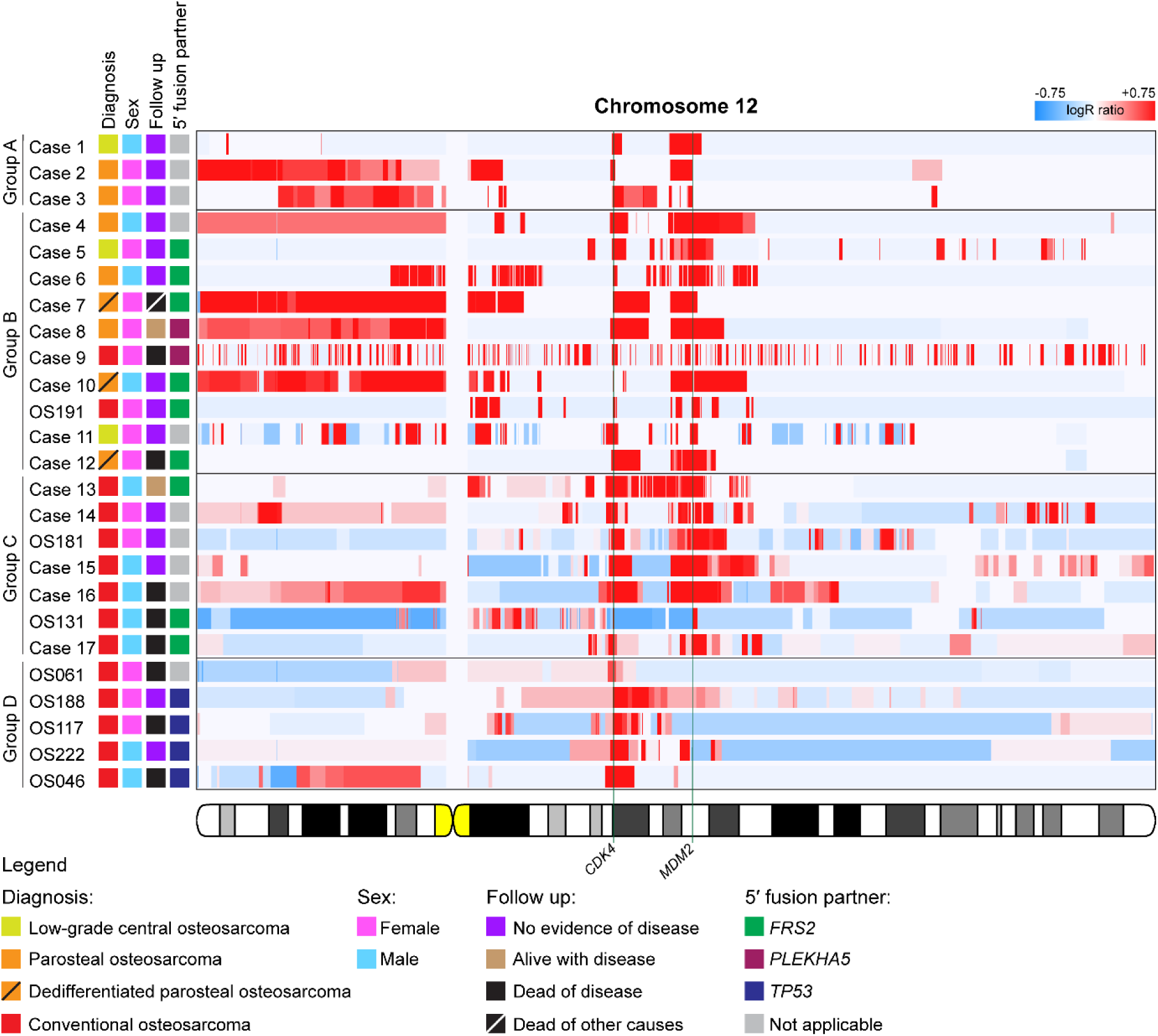
Chromosome 12 copy number heatmap of 25 osteosarcomas with amplification of 12q sequences. Copy number gains (depicted in red) and losses (depicted in blue) affecting chromosome 12 were identified by copy number array analysis. The cases are categorised into four groups based on their genome-wide copy number alterations and *MDM2* status. The loci of the *CDK4* and *MDM2* genes are indicated by a green line.

**Figure 3.**
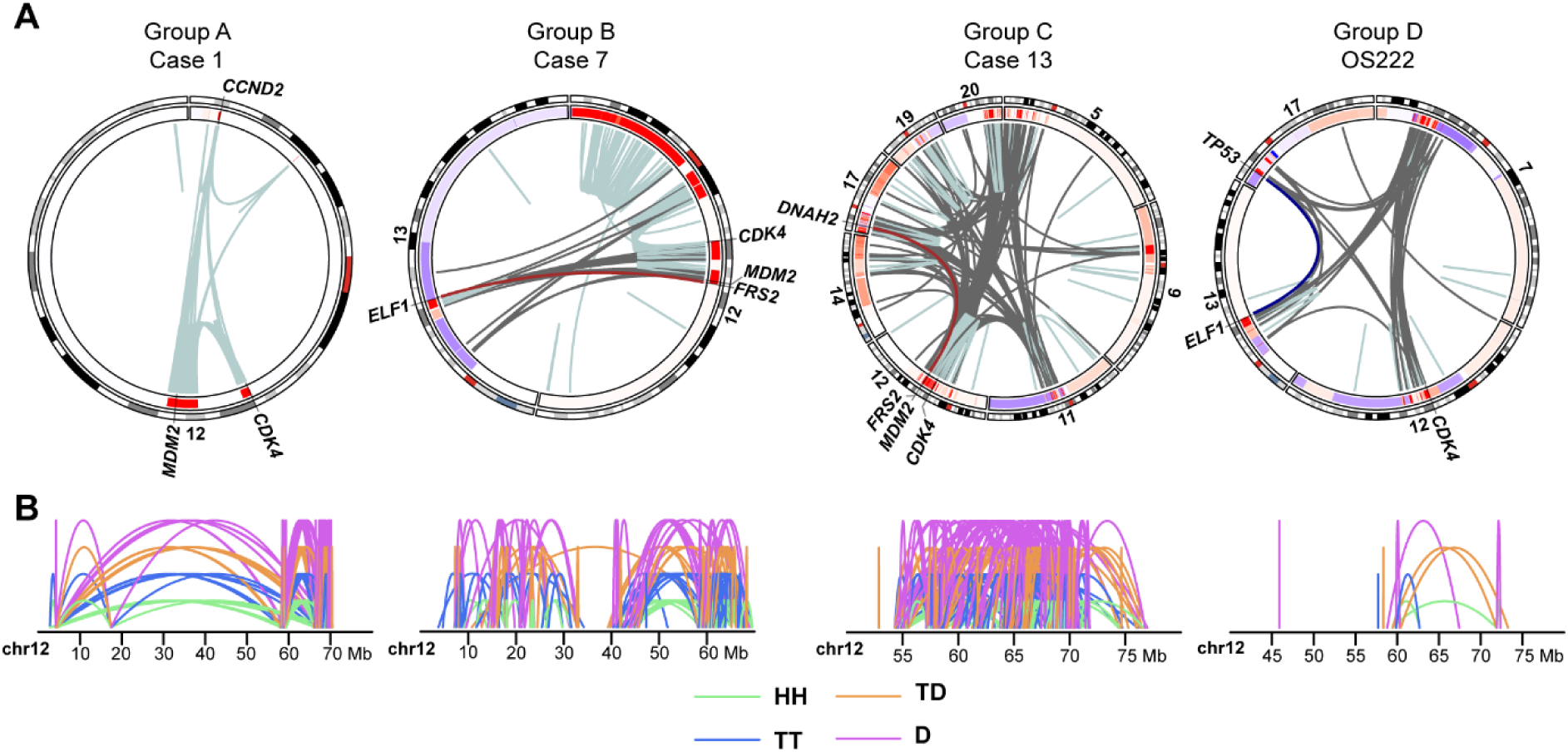
Combined copy number and structural variant data from representative cases. **(A)** Circos plots of representative cases from each subgroup are presented. In the circular track under the ideograms, red regions indicate copy number gains and amplifications, while blue regions indicate copy number loss. Intrachromosomal rearrangements are depicted in light blue and interchromosomal rearrangements in grey. Selected rearrangements of the *FRS2* and *TP53* promoter regions are displayed in brown and dark blue, respectively. **(B)** Intrachromosomal structural variants on chromosome 12 are depicted by mapping orientation of the read-pairs. Abbreviations: HH = head-to-head inversion, TT = tail-to-tail inversion, TD = duplication type and D = deletion type. Mb = mega-base-pair.

### Group B *CDK4* and *MDM2* amplified osteosarcomas with few alterations outside of chromosome 12

Group B included ten osteosarcomas, spanning low- and high-grade cases (three parosteal, two low-grade central, three dedifferentiated parosteal, and two conventional), all displaying amplification of *CDK4* and *MDM2,* along with regions on chromosome arm 12p in all but three cases (Figures 1 and 2, Supplementary Figure S2 and Supplementary Table S1). There were no deletions affecting chromosome 12, except in one case, and relatively few alterations beyond chromosome 12. Recurrent structural variants affected the *FRS2* (12q15) and *PLEKHA5* (12p12) genes. The 5′ parts of these genes, including their respective promoter region, underwent transposition, leading to elevated expression levels of other genes under their regulatory influence (Figure 4A-B, Supplementary Figure S2 and Supplementary Tables S2 and S3). Single-cell whole-genome analyses of three cases indicated the presence of coexisting subclones (Supplementary Figure S2). The distribution and read orientation of sequencing reads from chromosome 12 suggested a chromothripsis event (Figure 3, Supplementary Figure S2) [19]. In summary, Group B amplicons likely originated from an initial chromothripsis event affecting an extra copy of chromosome 12, followed by subsequent rounds of selective amplification. In some cases, parts of other chromosomes were intertwined with the chromosome 12 amplicons, leading to gene dysregulation through promoter swapping events.

**Figure 4.**
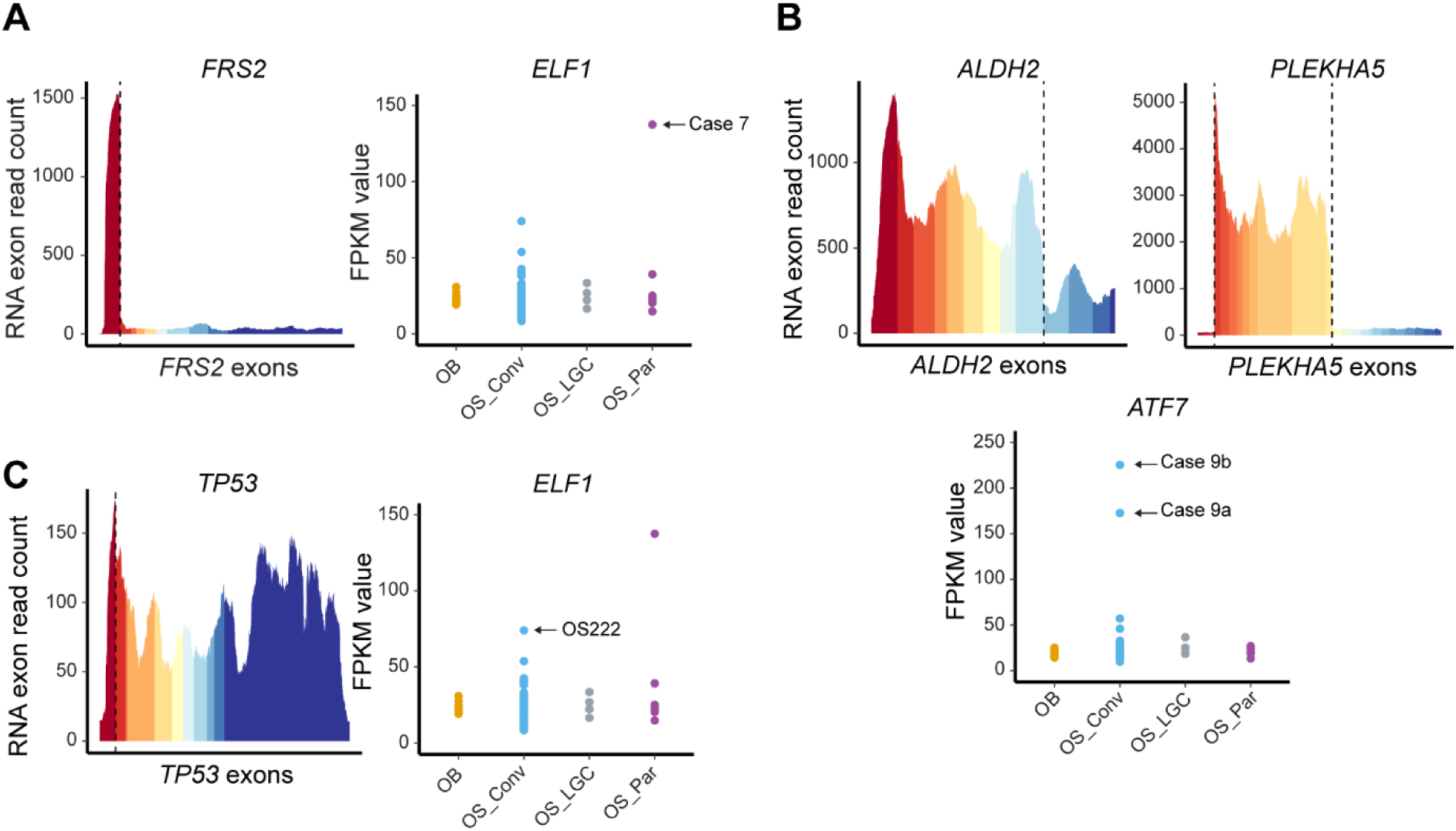
Promoter swapping events lead to gene upregulation. **(A)** In Case 7, the complete coding sequence of the *ELF1* gene is positioned under the control of the *FRS2* promoter. *FRS2* exon 1 displays a higher expression compared to the exons excluded from the fusion. *ELF1* is upregulated in Case 7 (indicated by an arrow) compared to other osteosarcomas and osteoblastomas. **(B)** Complex rearrangements in Case 9 lead to the formation of a three-way *ALDH2::PLEKHA5::ATF7* promoter swapping event. The *ALDH2* and *PLEKHA5* exons included in the fusion display a higher expression than the excluded exons. The *ATF7* gene in the multi-sampled Case 9 (indicated by two arrows) is upregulated compared to other osteosarcomas and osteoblastomas. **(C)** In OS222, the complete coding sequence of the *ELF1* gene is placed under the control of the *TP53* promoter. *TP53* exon 1 displays a higher expression than the exons excluded from the fusion. The *ELF1* gene in OS222 (indicated by an arrow) is highly expressed compared to other osteosarcomas and osteoblastomas, but lower than in Case 7. The RNA breakpoints are represented by dashed lines. Abbreviations: OB = osteoblastoma, OS_Conv = conventional osteosarcoma, OS_LGC = low-grade central osteosarcoma and OS_Par = parosteal or dedifferentiated parosteal osteosarcoma.

### Group C *CDK4* and *MDM2* amplified osteosarcomas with extensively altered genomes

Group C comprised seven conventional osteosarcomas displaying amplification of *CDK4* and *MDM2,* along with numerous genome-wide copy number and structural variants (Figures 1 and 2, Supplementary Figure S3 and Supplementary Table S1). Relative copy number loss of parts of chromosome 12 was observed in all but one case. The amplified material from chromosome 12 was intricately intertwined and co-amplified with material from several other chromosomes (Figure 3, Supplementary Figure S3). Structural variants transposing the *FRS2* promoter region were identified in three cases (Supplementary Table S2). The high genome complexity, coupled with limitations in detecting very high copy numbers, hindered a reliable evaluation of the potential presence of subclones via single-cell sequencing (Supplementary Figure S3). The distribution and read orientation of sequencing reads from chromosome 12 suggested a chromothripsis event in some cases in this group. In the remaining cases, there was inconclusive evidence of chromothripsis affecting chromosome 12, raising the possibility of other mechanisms being responsible for the initial onset of rearrangements and amplifications in these cases (Figure 3, Supplementary Figure S3) [19]. In summary, it is likely that some Group C amplicons originated from an initial chromothripsis event affecting either an extra copy or one of the normal chromosome 12 homologues, followed by subsequent rounds of selective amplification. Promoter swapping events affecting the *FRS2* gene were detected in both Group B and C.

### Group D *CDK4*-amplified and *TP53*-rearranged osteosarcomas with extensively altered genomes

Group D consisted of five conventional osteosarcomas exhibiting amplification of *CDK4*, rearrangements of *TP53,* and numerous genome-wide copy number and structural variants (Figures 1 and 2, Supplementary Figure S4 and Supplementary Table S1). In all cases, deletions affecting chromosome 12 were observed. The amplified material from chromosome 12 was intricately intertwined and co-amplified with material from several other chromosomes (Figure 3, Supplementary Figure S4). One case harboured a homozygous deletion of *TP53*, while the remaining four cases displayed structural variants transposing the *TP53* promoter region. The active *TP53* promoter was fused to and upregulated various 3′ partner genes [17] (Figure 4C, Supplementary Figure S4 and Supplementary Table S1). There was no evidence of a chromothripsis event affecting chromosome 12 in these cases; instead, the initial disruption of the *TP53* locus led to the creation of a dicentric chromosome. Subsequent break and repair processes intertwined and co-amplified the *CDK4* locus with the *TP53* promoter gene fusion, along with parts of other chromosomes involved. In the case with homozygous loss of *TP53*, no clear connection was identified between rearrangements around the lost *TP53* locus and *CDK4* amplification.

### Osteosarcomas exhibiting *CDK4* amplification comprise of distinct genetic subgroups

All cases presented amplification of *CDK4*. Groups A, B and C exhibited concurrent amplification of *MDM2* in increasingly complex genomes, whereas Group D harboured *TP53* promoter gene fusions or *TP53* homozygous loss. For all cases in Groups A and B and most in Group C, an initial chromothripsis event affecting either an extra copy or one of the normal homologues of chromosome 12 was detected. This was verified for selected cases using longread whole-genome sequencing where the same patterns of copy number amplification and clustering of structural variants were seen as in the initial analysis (Supplementary Figure S5).

Analyses of global gene expression levels revealed that *TP53*-wildtype tumours clustered separately from *TP53*-mutated tumours (Figure 5A). This was most evident in cases from Group C, where *TP53*-wildtype Case 15 clustered with Group B, while Case 17 harbouring two single nucleotide variants in *TP53* clustered with Group D. In this dataset, *HMGA2* amplification and *TP53* mutation were observed as mutually exclusive events (Supplementary Table S1 and Supplementary Figure S6). Partial or full amplification of the *HMGA2* gene was observed in all cases within Groups A, B and C, except for OS131 and Case 17 in Group C, both of which exhibit concurrent *MDM2* amplification and *TP53* mutation. Notably, *HMGA2* amplification was not detected in Group D.

**Figure 5.**
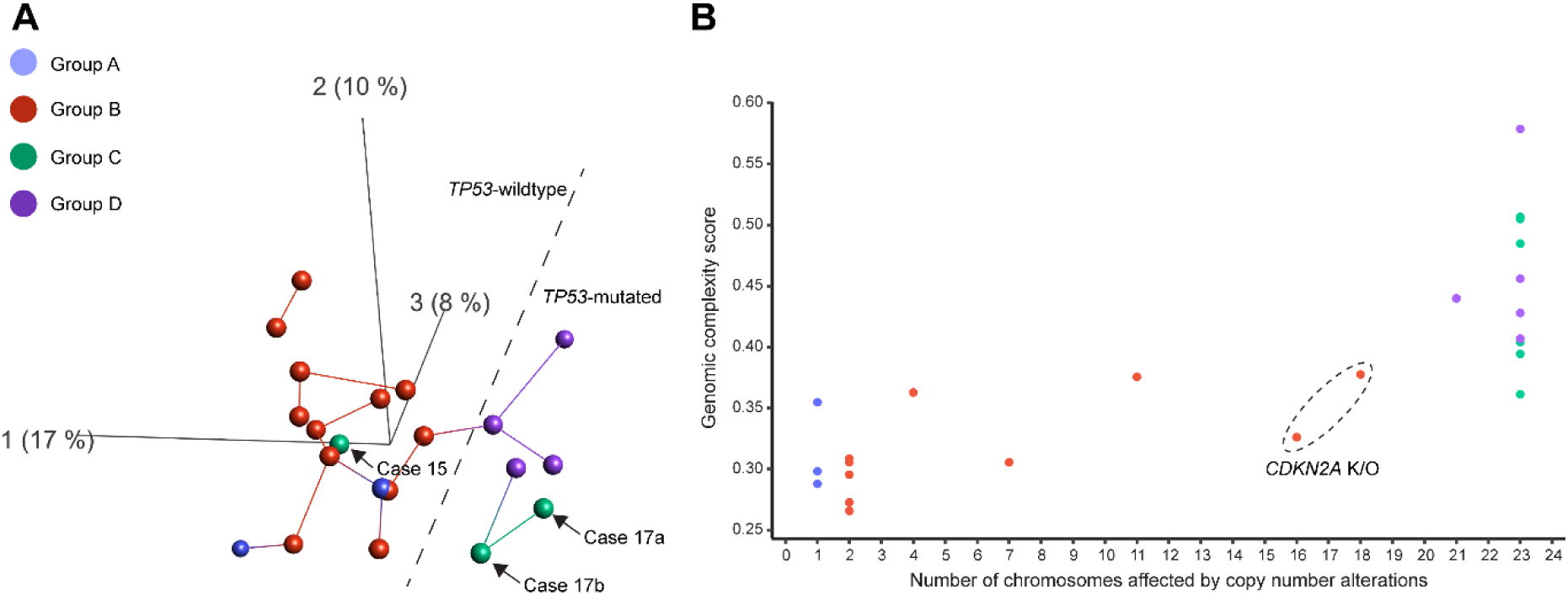
*CDK4*-amplified osteosarcomas show distinct gene expression profiles and diverse genomic complexity levels. **(A)** Unsupervised principal component analyses based on global gene expression levels in osteosarcomas reveal distinct profiles. Cases from Groups B and D are separated in two distinct clusters. However, cases from Group C cluster with both Group B and D. A notable difference in global gene expression is instead determined by the status of the *TP53* gene, where mutated cases cluster distinctly from wildtype cases. The first three principal components representing 17%, 10% and 8% of the variation are displayed. Each sample is connected with its nearest neighbour. Samples to the left of the dashed line are *TP53*-wildtype, while those to the right are *TP53*-mutated. **(B)** The genomic complexity score of each case plotted against the number of chromosomes affected by copy number alterations. Cases from Groups A and B formed one cluster with relatively few chromosomes affected by copy number alterations and lower genomic complexity, while those from Groups C and D formed another cluster with virtually every chromosome affected and relatively high genomic complexity. Cases 7 and 8 from Group B (enclosed within a dashed ellipse), both with complete knock-out of *CDKN2A*, showed copy number alterations on more chromosomes than other cases in Group B. A chromosome was counted as being affected by a copy number alteration if there was a visible copy number shift in the segmentation analysis, or if the chromosome displayed a non-diploid copy number. The X and Y chromosomes were counted as one chromosome pair in males.

Moreover, a genomic complexity score (GCS) was calculated for each case based on SNP array copy number data and plotted against the number of chromosomes exhibiting copy number alterations [23]. This analysis revealed two primary clusters among the cases: one characterised by relatively low genetic complexity, with 11 or fewer chromosomes affected by copy number alterations (except for two cases with homozygous loss of *CDKN2A*); and another with higher genetic complexity, involving nearly all chromosomes displaying copy number alterations (Figure 5B and Supplementary Table S1). The first cluster included cases assigned to Groups A and B, initially distinguished by whether only chromosome 12 was affected by copy number alterations or if other chromosomes were also affected. The second cluster comprised cases from Groups C and D, exhibiting highly altered genomes and further subdivided based on the presence of either *MDM2* amplification or *TP53* alterations.

In Groups B and C, recurrent rearrangements were observed in the *PLEKHA5* and *FRS2* genes, similar to the *TP53* promoter rearrangements seen in Group D. Specifically, Cases 8 and 9 harboured the *PLEKHA5::EPS8* and *ALDH2::PLEKHA5::ATF7* fusions, respectively, with a 30 kb region of *PLEKHA5* shared between the two fusions at the DNA level (Figure 4B, Supplementary Figures S2 and Supplementary Tables S2 and S3). The functional significance of this shared sequence for the respective *PLEKHA5* fusion remains unclear. Notably, both cases with *PLEKHA5* rearrangements showed a particularly aggressive clinical behaviour (Supplementary Table S1). Among the *FRS2*-rearranged cases classified as conventional osteosarcoma, the consequences of a dislocated *FRS2* promoter could not be determined. The remaining five *FRS2*-rearranged cases were low-grade osteosarcomas or of low-grade origin. All *FRS2* promoter partner genes, with one exception, demonstrated the highest expression levels in the affected cases compared with other osteosarcomas (Figure 4A and Supplementary Figure S2). In cases with *PLEKHA5* and *FRS2* rearrangements that were multi-sampled, the respective fusions were present in all investigated tumour materials (core needle biopsy, resection specimen, recurrence and/or metastasis), except for Case 12 (*FRS2::ADAM32*). In this case, the fusion was detected in the primary tumour resection but not in the metastatic sample taken almost 20 years later, potentially indicating a slower growth rate without the fusion (Supplementary Tables S1 and S2 and Supplementary Figure S5). It is worth noting that one 3′ fusion partner was recurrent in the present study. The *ELF1* gene was upregulated by both the *FRS2* and *TP53* promoters, respectively, resulting in the highest and second highest *ELF1* expression among all 95 RNA-sequenced samples (Figure 4A,C and Supplementary Figures S2D and S4B).

## Discussion

We demonstrate that genetic rearrangements in osteosarcoma result in ectopic gene expression through two mechanisms: gene amplification and transposition of regulatory elements. Our data support that these events are intertwined and ongoing, at least during tumour initiation. This process facilitates the emergence of subclones with varying fitness, creating a substrate for selection during tumour evolution. In low-grade cases and those with a low-grade origin, this diverse subclonal architecture persists and is detectable at the single cell level – a typical finding previously identified through G-banding analysis [8, 24]. Given the potential dedifferentiation of low-grade cases, this emergence of subclones is anticipated. However, in the high-grade cases, such phenomena are more challenging to detect due to the substantial number and magnitude of alterations.

Selective amplification of *CDK4* and *MDM2* is a particularly common phenomenon in low-grade osteosarcoma. However, in high-grade cases, such amplifications are far less frequent [8–11]. Within our unselected cohort of high-grade osteosarcomas, 5% exhibited amplification of *CDK4* alone, and only 3% displayed amplification of both *CDK4* and *MDM2*. It is noteworthy that these amplification patterns are somewhat mirrored in other tumour types. For instance, ring chromosomes containing amplified copies of *MDM2*, often accompanied by *CDK4*, are also present in subtypes of soft tissue tumours. They represent the genetic hallmark of well-differentiated liposarcoma and its dedifferentiated counterpart [9, 25–28]. Recently, Sydow et al. suggested that amplification of sequences from chromosome 12 in lipomatous tumours does not occur through classical chromothripsis [19, 29]. Instead, they argue that large segments are most often gained after DNA synthesis and that these segments can oscillate between circularized and rod-shaped configurations. When circularized, amplification of selected segments is achieved through repeated breakage-fusion-bridge cycles. Moreover, co-amplification of segments from other chromosomes often appears to occur as a secondary event.

In osteosarcoma, the most biologically significant alteration appears to be the presence of *TP53* mutation, manifested either as structural changes or single nucleotide variants [14–18, 30]. In cases with *TP53* structural variants, previous reports indicate that this event occurs early, likely serving as the initiating event in at least half of paediatric osteosarcomas [15, 17]. Here, we illustrate that a subset of the *TP53* promoter fusion-positive cases co-amplify *CDK4* during this early process. Unlike *MDM2*-amplified osteosarcomas, *TP53*-rearranged cases show no evidence of chromothripsis affecting chromosome 12. These cases do not seem to benefit from increased copy numbers of *MDM2*, suggesting that altered *TP53* function takes precedence over *MDM2* amplification [17].

Our recent publication on *TP53* structural alterations in osteosarcoma did not reveal global gene expression patterns dependent on *TP53* status [17]. However, in the current cohort, *TP53*-mutated osteosarcomas formed a distinct cluster separate from *TP53*-wildtype tumours. We attribute this difference to the fact that the former analysis encompassed a much more heterogeneous cohort [17, 23]. In the current analysis, the relative contribution of an altered *TP53* pathway was not clouded by a great diversity in other genetic features of the investigated tumours. Furthermore, the tumour biology differs between *TP53*-rearranged and *MDM2*-amplified osteosarcomas. Generally, *TP53*-rearranged tumours are predominantly found in children and adolescents [17], while *MDM2* amplification is more frequently detected in osteosarcomas of young adults and adults. This difference suggests that *MDM2*-amplified osteosarcoma have distinct origins and biological characteristics compared to *TP53*-rearranged osteosarcomas.

The backbone of detected amplicons in *MDM2*-amplified osteosarcomas consists mostly of chromosome arm 12q sequences, likely arising from a chromothripsis event affecting chromosome 12. Subsequent rounds of breakage-fusion-bridge cycles lead to copy number amplification [20–22]. In some cases, material from other chromosomes is intertwined and co-amplified through such cycles or possibly through punctuated chromothripsis episodes. Among the co-amplified sequences were the recurrent *FRS2* and *PLEKHA5* gene fusions. Co-amplification of *FRS2* and *MDM2* in low-grade and dedifferentiated osteosarcoma has been previously reported [31]. However, these findings were based on FISH analysis using a probe located 5′ of the *FRS2* gene, making it difficult to assess whether whole or only parts of *FRS2* were amplified. This leaves the possibility open for the amplified copies to be rearranged, leading to *FRS2* fusions similar to what we describe here. We do not have evidence that unequivocally supports a functional role for the *FRS2* and *PLEKHA5* fusion events, and further studies are warranted to determine their role in tumour progression. Supporting a possible biological significance is the fact that these events are recurrent, induce the expression of the 3′ partner genes, and cluster in *MDM2*-amplified osteosarcomas with either aggressive clinical behaviour or dedifferentiated morphology. It should also be noted that the high complexity of these recurrent genetic structures supports, rather than rules out, functional importance, although their function may be more challenging to unravel. Perhaps the most intriguing finding in this context is the fact that the *ELF1* gene was upregulated through fusion to both the *TP53* and the *FRS2* promoter, respectively.

In conclusion, our study reveals that osteosarcomas with *CDK4* amplification belong to distinct genetic subgroups characterised by recurrent patterns of additional alterations. These patterns confer biological consequences that manifest as clinically detectable differences, including variations in subtype, age of onset, and outcome.

## Materials and Methods

### Tumour material and clinical features

The present study encompassed a total of 25 selected cases, with frozen tissue available from more than one time point and/or location for seven of them. The samples comprised diagnostic specimens, primary tumour resections, as well as local and distant recurrences, with the most long-term being obtained 18 years after the initial diagnosis. Additional clinical information on the selected cases is provided in Supplementary Table S1.

### Ethics statement

All tumour material was obtained after informed consent, and the study was approved by the Swedish Ethics Review Authority (Dnr 2023-01550-01) and the Ethikkommission beider Basel (reference 274/12).

### Genome-wide DNA copy number analyses

DNA was extracted according to standard procedures from fresh frozen tumour biopsies and hybridized to CytoScan HD arrays, following protocols supplied by the manufacturer (Thermo Fisher Scientific, Waltham, MA, USA). Data analysis was performed using the Chromosome Analysis Suite v 4.1.0.90 (Thermo Fisher Scientific), detecting imbalances by visual inspection, and by segmenting log_2_ values using the R packages copynumber [32] and TAPS [33]. Somatic copy number alterations, based on SNP array data or technologies with lower resolution, have been published previously for a subset of cases (Supplementary Table S1). A genomic complexity score (GCS) was calculated for each case to reflect the degree of genomic complexity based on copy number alterations as previously described [23].

### Whole-genome mate pair sequencing for detection of structural aberrations

For the detection of structural chromosomal abnormalities, mate pair libraries were prepared for sequencing using the Nextera mate pair sample preparation kit (Illumina, San Diego, CA, USA), as previously described [34]. Sequencing depth was on average 2.72x (mapping coverage 1.96x) and the mean insert size was 3.0 kb, resulting in a median spanning coverage of 53.2x of the human genome (mean 53.0x, range 11.4x-119.1x). All samples were sequenced with high quality and yield; between 13.1 and 115.5 million read pairs were obtained per sample (mean 53.8 million read pairs) and the average quality scores were 31.5-33.5. Sequencing reads were trimmed using NxTrim v 0.4.2 [35] and subsequently aligned against the GRCh37/hg19 build using the Borrows-Wheeler Aligner v 0.7.15 [36]. To identify structural rearrangements, the sequence data were analysed using Integrative Genomics Viewer [37, 38], as well as the structural variant callers TIDDIT v 2.12.1, Delly2 v 0.7.8 and Manta v 1.2.2 [39–41]. Structural alterations were considered true when identified by at least two of the three variant callers. Copy number and structural variant data were then combined to produce circos plots using the R package ‘circlize’ [42] as previously described [17].

### Whole-genome longread sequencing for verification of structural alterations

For the verification of structural chromosomal abnormalities detected with mate pair whole-genome sequencing, selected samples where high molecular weight DNA could be obtained were subjected to longread whole-genome sequencing on the Pacific Biosciences Revio Technology Platform, using the HiFi protocol (PacBio, Menlo Park, CA, USA). DNA was extracted using the PacBio Nanobind Tissue Kit according to the manufacturer’s instructions. Raw sequencing data was aligned to the GRCh38/hg38 build of the human reference genome using pbmm2 as part of SMRTLink (PacBio) v 13 and structural variants were detected using pbsv. Due to a lack of normal controls, only structural variants annotated as “BND” or “INV” and passing all quality filters were considered for further analyses. Coverage levels were calculated using Mosdepth v 0.3.6 [43] with a bin size of 100000 bp. Coverage levels and structural variant data were then combined to produce circos plots using the R package ‘circlize’ [42] as previously described [17].

### Whole-genome low-pass sequencing of single cells

Whole-genome sequencing of cryopreserved primary osteosarcoma cells was performed as described in detail previously [44]. In brief, library preparation was performed using a modified single-cell whole-genome sequencing protocol and 77 base pair single reads were generated using a NextSeq 500 sequencing instrument (Illumina). From each assessed case, either 48 or 96 single cells were subjected to sequencing at an average depth of 0.01x and between 29-82 single cells per assessed case passed quality control. This included varying amounts of normal non-neoplastic cells. Copy number analysis was performed using AneuFinder [45].

### RNA sequencing for detection of gene fusions, expression levels, single nucleotide variants and indels

RNA was extracted according to standard procedures from fresh frozen tumour biopsies and sequenced as previously described [34]. FusionCatcher v 1.0 and STAR-Fusion v 1.4.0 were used to identify candidate fusion transcripts from the sequence data [46, 47]. Fusions were considered noise if identified by only one of the programs, or if there was no support for the fusion in whole-genome mate pair sequencing data. NAFuse was used for automatic matching of RNA and DNA sequencing data for gene fusion verification [23]. Sequencing reads were aligned to the GRCh37/hg19 build using STAR v 2.5.2b [48]. For comparison of relative gene expression levels, transcript quantification and normalisation was carried out with RSEM [49]. Data was visualised using the Qlucore Omics Explorer version 3.8 (Qlucore AB, Lund, Sweden). Relative gene expression levels were evaluated in conventional (*n* = 69), dedifferentiated parosteal (*n* = 3), parosteal (*n* = 4), and low-grade central (*n* = 3) osteosarcomas, as well as osteoblastomas (*n* = 13). For three 12q-amplified samples, RNA-sequencing data from multiple samples was included. Single nucleotide variants and indels were detected using VarScan v 2.4.1, MuTect v 1.1.7 and Mutect2 [50–52]. Constitutional variants were excluded based on information from the Genome Aggregation Database (gnomAD v2.1.1) [53]. The detected variants were finally confirmed by manual inspection using the Integrative Genomics Viewer.

### Fluorescence in situ hybridization

Fluorescence in situ hybridization (FISH) analyses on metaphase spreads were performed as described previously [54], using in-house labelled bacterial artificial chromosome clones or commercially available probes specific for the *FRS2* (RP11-1102B16), *CDK4* (RP11-571M6), and *MDM2* (Vysis LSI MDM2) genes, and whole chromosome painting probes (BACPAC Genomics, Emeryville, CA, USA, Applied Spectral Imaging, Carlsbad, CA, USA and Abbott Molecular, Abbott Park, IL, USA).

## Data availability

Sequencing data for a subset of cases had been previously deposited at the European Genome-Phenome Archive (EGA) under the accession number EGAS00001003842 [17].

Remaining data will also be uploaded to the EGA and an additional accession number will be made available.

## Acknowledgement

KHS was supported by the Royal Physiographic Society (Lund, Sweden). KHN was supported by the Swedish Childhood Cancer Fund, the Swedish Cancer Society, the Swedish Research Council, the Swedish state under the agreement between the Swedish government and the country councils (the ALF agreement), the Faculty of Medicine at Lund University, the Åke Wiberg Foundation, the Royal Physiographic Society (Lund, Sweden), and the Crafoord Foundation. DB was supported by the Foundation of the Basel Bone Tumour Reference Centre, the Gertrude von Meissner Stiftung, the Stiftung für krebskranke Kinder, Regio Basiliensis and the Basel Research Centre for Child Health (BRCCH). MN was supported by the Helga und Heinrich Holzhauerstiftung. The authors would like to acknowledge support of the National Genomics Infrastructure (NGI) / Uppsala Genome Center and UPPMAX for providing assistance in massive parallel sequencing and computational infrastructure. Work performed at NGI / Uppsala Genome Center has been funded by RFI / VR and Science for Life Laboratory, Sweden.

## Author contributions

KHS and KHN conceived and designed the experiments. ES, MN, FHdF, and DB contributed tumour material and clinical information. FHdF and DB performed histopathological review. KHS, VD, JN, LM, and KHN performed experimental and bioinformatic analyses, and interpreted the results. HvdB, DCJS and FF applied low-pass whole-genome sequencing on single cells. KHS, VD and KHN prepared the manuscript with contributions from all other authors.

## References

1. Gianferante DM, Mirabello L, Savage SA. Germline and somatic genetics of osteosarcoma - connecting aetiology, biology and therapy. Nat Rev Endocrinol 2017; 13: 480–491.

2. Baumhoer D, Böhling TO, Cates JMM, et al. Osteosarcoma. In: WHO Classification of Tumours Editorial Board. Soft Tissue and Bone Tumours. 5^th^ ed. International Agency for Research on Cancer; 2020: 403–409.

3. Beird HC, Bielack SS, Flanagan AM, et al. Osteosarcoma. Nat Rev Dis Primers 2022; 8: 77.

4. Yoshida A, Bredella MA, Gambarotti M, et al. Low-grade central osteosarcoma. In: WHO Classification of Tumours Editorial Board. Soft Tissue and Bone Tumours. 5^th^ ed. International Agency for Research on Cancer; 2020: 400–402.

5. Wang J, Nord KH, O’Donnell PG, et al. Parosteal osteosarcoma. In: WHO Classification of Tumours Editorial Board. Soft Tissue and Bone Tumours. 5^th^ ed. International Agency for Research on Cancer; 2020: 410–413.

6. Bertoni F, Bacchini P, Staals EL, et al. Dedifferentiated parosteal osteosarcoma: the experience of the Rizzoli Institute. Cancer 2005; 103: 2373–2382.

7. Choong PF, Pritchard DJ, Rock MG, et al. Low grade central osteogenic sarcoma. A long-term followup of 20 patients. Clin Orthop Relat Res 1996: 198–206.

8. Gisselsson D, Pålsson E, Höglund M, et al. Differentially amplified chromosome 12 sequences in low- and high-grade osteosarcoma. Genes Chromosomes Cancer 2002; 33: 133–140.

9. Heidenblad M, Hallor KH, Staaf J, et al. Genomic profiling of bone and soft tissue tumors with supernumerary ring chromosomes using tiling resolution bacterial artificial chromosome microarrays. Oncogene 2006; 25: 7106–7116.

10. Szymanska J, Mandahl N, Mertens F, et al. Ring chromosomes in parosteal osteosarcoma contain sequences from 12q13-15: a combined cytogenetic and comparative genomic hybridization study. Genes Chromosomes Cancer 1996; 16: 31–34.

11. Mejia-Guerrero S, Quejada M, Gokgoz N, et al. Characterization of the 12q15 *MDM2* and 12q13-14 CDK4 amplicons and clinical correlations in osteosarcoma. Genes Chromosomes Cancer 2010; 49: 518–525.

12. Dujardin F, Binh MB, Bouvier C, et al. *MDM2* and *CDK4* immunohistochemistry is a valuable tool in the differential diagnosis of low-grade osteosarcomas and other primary fibro-osseous lesions of the bone. Mod Pathol 2011; 24: 624–637.

13. Salinas-Souza C, De Andrea C, Bihl M, et al. *GNAS* mutations are not detected in parosteal and low-grade central osteosarcomas. Mod Pathol 2015; 28: 1336–1342.

14. Behjati S, Tarpey PS, Haase K, et al. Recurrent mutation of *IGF* signalling genes and distinct patterns of genomic rearrangement in osteosarcoma. Nat Commun 2017; 8: 15936.

15. Chen X, Bahrami A, Pappo A, et al. Recurrent somatic structural variations contribute to tumorigenesis in pediatric osteosarcoma. Cell Rep 2014; 7: 104–112.

16. Lorenz S, Barøy T, Sun J, et al. Unscrambling the genomic chaos of osteosarcoma reveals extensive transcript fusion, recurrent rearrangements and frequent novel TP53 aberrations. Oncotarget 2016; 7: 5273–5288.

17. Saba KH, Difilippo V, Kovac M, et al. Disruption of the *TP53* locus in osteosarcoma leads to *TP53* promoter gene fusions and restoration of parts of the *TP53* signalling pathway. J Pathol 2024; 262: 147–160.

18. Smida J, Xu H, Zhang Y, et al. Genome-wide analysis of somatic copy number alterations and chromosomal breakages in osteosarcoma. Int J Cancer 2017; 141: 816–828.

19. Korbel JO, Campbell PJ. Criteria for inference of chromothripsis in cancer genomes. Cell 2013; 152: 1226–1236.

20. Garsed DW, Marshall OJ, Corbin VD, et al. The architecture and evolution of cancer neochromosomes. Cancer Cell 2014; 26: 653–667.

21. Gisselsson D, Pettersson L, Höglund M, et al. Chromosomal breakage-fusion-bridge events cause genetic intratumor heterogeneity. Proc Natl Acad Sci U S A 2000; 97: 5357–5362.

22. Shoshani O, Brunner SF, Yaeger R, et al. Chromothripsis drives the evolution of gene amplification in cancer. Nature 2021; 591: 137–141.

23. Difilippo V, Saba KH, Styring E, et al. Osteosarcomas With Few Chromosomal Alterations or Adult Onset Are Genetically Heterogeneous. Lab Invest 2024; 104: 100283.

24. Mertens F, Mandahl N, Orndal C, et al. Cytogenetic findings in 33 osteosarcomas. Int J Cancer 1993; 55: 44–50.

25. Nord KH, Macchia G, Tayebwa J, et al. Integrative genome and transcriptome analyses reveal two distinct types of ring chromosome in soft tissue sarcomas. Hum Mol Genet 2014; 23: 878–888.

26. Pedeutour F, Forus A, Coindre JM, et al. Structure of the supernumerary ring and giant rod chromosomes in adipose tissue tumors. Genes Chromosomes Cancer 1999; 24: 30–41.

27. Pedeutour F, Suijkerbuijk RF, Forus A, et al. Complex composition and co-amplification of SAS and MDM2 in ring and giant rod marker chromosomes in well-differentiated liposarcoma. Genes Chromosomes Cancer 1994; 10: 85–94.

28. Wang X, Asmann YW, Erickson-Johnson MR, et al. High-resolution genomic mapping reveals consistent amplification of the fibroblast growth factor receptor substrate 2 gene in well-differentiated and dedifferentiated liposarcoma. Genes Chromosomes Cancer 2011; 50: 849–858.

29. Sydow S, Piccinelli P, Mitra S, et al. MDM2 amplification in rod-shaped chromosomes provides clues to early stages in circularized gene amplification in liposarcoma. Submitted manuscript 2023.

30. Ribi S, Baumhoer D, Lee K, et al. TP53 intron 1 hotspot rearrangements are specific to sporadic osteosarcoma and can cause Li-Fraumeni syndrome. Oncotarget 2015; 6: 7727–7740.

31. He X, Pang Z, Zhang X, et al. Consistent Amplification of FRS2 and MDM2 in Low-grade Osteosarcoma: A Genetic Study of 22 Cases With Clinicopathologic Analysis. Am J Surg Pathol 2018; 42: 1143–1155.

32. Nilsen G, Liestøl K, Van Loo P, et al. Copynumber: Efficient algorithms for single- and multi-track copy number segmentation. BMC Genomics 2012; 13: 591.

33. Rasmussen M, Sundström M, Göransson Kultima H, et al. Allele-specific copy number analysis of tumor samples with aneuploidy and tumor heterogeneity. Genome Biol 2011; 12: R108.

34. Saba KH, Cornmark L, Rissler M, et al. Genetic profiling of a chondroblastoma-like osteosarcoma/malignant phosphaturic mesenchymal tumor of bone reveals a homozygous deletion of *CDKN2A*, intragenic deletion of *DMD*, and a targetable *FN1-FGFR1* gene fusion. Genes Chromosomes Cancer 2019; 58: 731–736.

35. O’Connell J, Schulz-Trieglaff O, Carlson E, et al. NxTrim: optimized trimming of Illumina mate pair reads. Bioinformatics 2015; 31: 2035–2037.

36. Li H. Toward better understanding of artifacts in variant calling from high-coverage samples. Bioinformatics 2014; 30: 2843–2851.

37. Robinson JT, Thorvaldsdóttir H, Winckler W, et al. Integrative genomics viewer. Nat Biotechnol 2011; 29: 24–26.

38. Thorvaldsdóttir H, Robinson JT, Mesirov JP. Integrative Genomics Viewer (IGV): high-performance genomics data visualization and exploration. Brief Bioinform 2013; 14: 178–192.

39. Chen X, Schulz-Trieglaff O, Shaw R, et al. Manta: rapid detection of structural variants and indels for germline and cancer sequencing applications. Bioinformatics 2016; 32: 1220–1222.

40. Eisfeldt J, Vezzi F, Olason P, et al. TIDDIT, an efficient and comprehensive structural variant caller for massive parallel sequencing data. F1000Research 2017; 6: 664.

41. Rausch T, Zichner T, Schlattl A, et al. DELLY: structural variant discovery by integrated paired-end and split-read analysis. Bioinformatics 2012; 28: i333–i339.

42. Gu Z, Gu L, Eils R, et al. circlize Implements and enhances circular visualization in R. Bioinformatics 2014; 30: 2811–2812.

43. Pedersen BS, Quinlan AR. Mosdepth: quick coverage calculation for genomes and exomes. Bioinformatics 2018; 34: 867–868.

44. van den Bos H, Bakker B, Taudt A, et al. Quantification of Aneuploidy in Mammalian Systems. Methods Mol Biol 2019; 1896: 159–190.

45. Bakker B, Taudt A, Belderbos ME, et al. Single-cell sequencing reveals karyotype heterogeneity in murine and human malignancies. Genome Biol 2016; 17: 115.

46. Haas BJ, Dobin A, Li B, et al. Accuracy assessment of fusion transcript detection via read-mapping and de novo fusion transcript assembly-based methods. Genome Biol 2019; 20: 213–213.

47. Nicorici D, Satalan M, Edgren H, et al. FusionCatcher – a tool for finding somatic fusion genes in paired-end RNA-sequencing data. bioRxiv 2014: 011650.

48. Dobin A, Davis CA, Schlesinger F, et al. STAR: ultrafast universal RNA-seq aligner. Bioinformatics 2013; 29: 15–21.

49. Li B, Dewey CN. RSEM: accurate transcript quantification from RNA-Seq data with or without a reference genome. BMC Bioinformatics 2011; 12: 323.

50. Benjamin D, Sato T, Cibulskis K, et al. Calling Somatic SNVs and Indels with Mutect2. bioRxiv 2019: 861054.

51. Cibulskis K, Lawrence MS, Carter SL, et al. Sensitive detection of somatic point mutations in impure and heterogeneous cancer samples. Nat Biotechnol 2013; 31: 213–219.

52. Koboldt DC, Zhang Q, Larson DE, et al. VarScan 2: somatic mutation and copy number alteration discovery in cancer by exome sequencing. Genome Res 2012; 22: 568–576.

53. Karczewski KJ, Francioli LC, Tiao G, et al. The mutational constraint spectrum quantified from variation in 141,456 humans. Nature 2020; 581: 434–443.

54. Jin Y, Möller E, Nord KH, et al. Fusion of the *AHRR* and *NCOA2* genes through a recurrent translocation t(5;8)(p15;q13) in soft tissue angiofibroma results in upregulation of aryl hydrocarbon receptor target genes. Genes Chromosomes Cancer 2012; 51: 510–520.

