## Supplementary Figures S1-S7 for "*CDK4* is co-amplified with either *TP53* promoter gene fusions or *MDM2* through distinct mechanisms in osteosarcoma"

Figure S1

#### A Case 1 - Low-grade central osteosarcoma

circos plot:

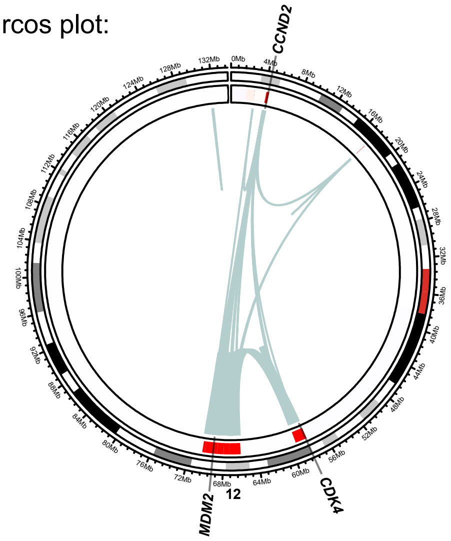


structural variant distribution:

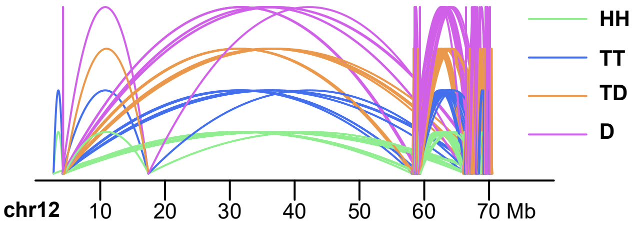


#### B Case 2 - Parosteal osteosarcoma

circos plot:

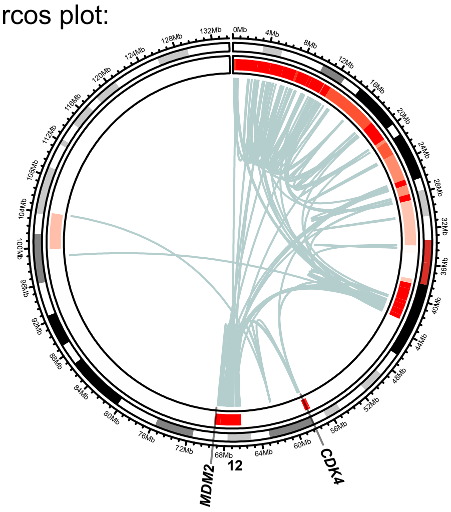


structural variant distribution:

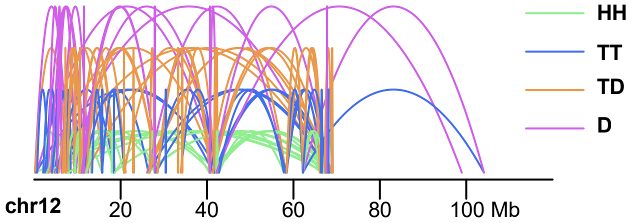


single cell whole-genome heatmap:

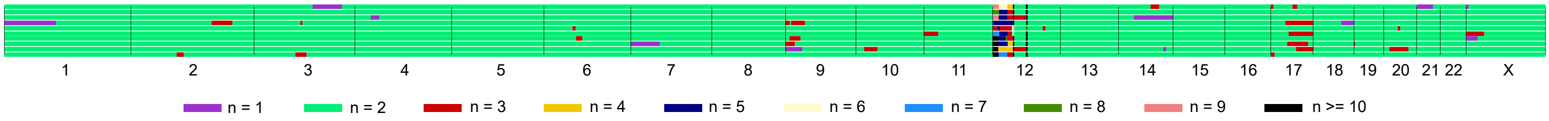


representative single cell:

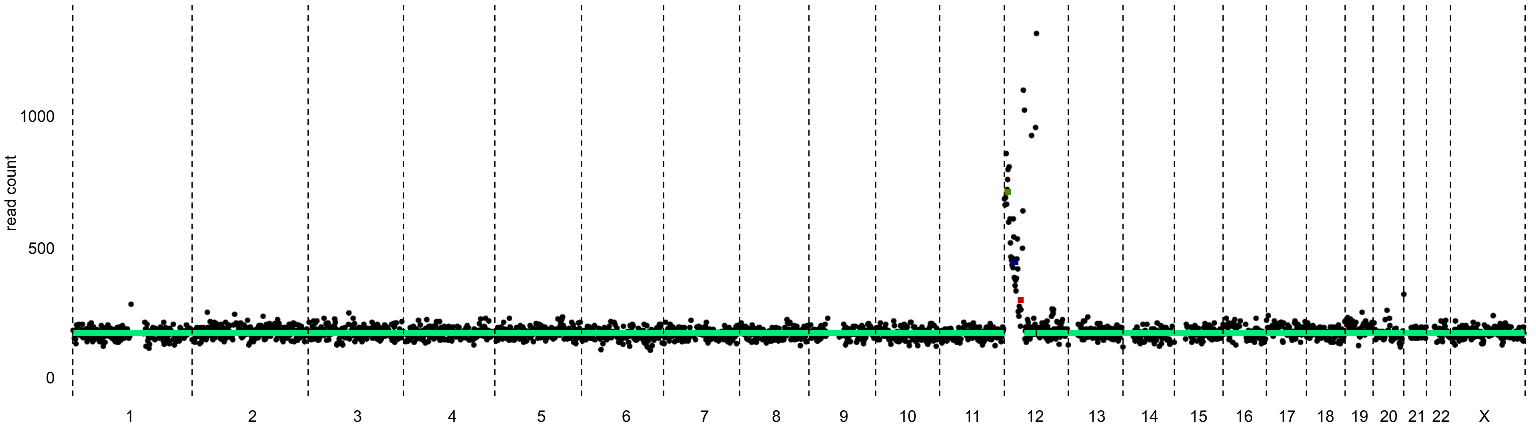

**Figure S1. Group A: combined copy number and structural variant data. Circos plot:** Red regions in the inner circular track of the circos plot indicate copy number gains, with higher level amplifications being in a more intense shade and lower-level gains in a lighter shade. Intrachromosomal structural variants are depicted in light blue. **Structural variant distribution:** Intrachromosomal structural variants plotted based on read mapping orientation. Abbreviations: HH = head-to-head inversion, TT = tail-to-tail inversion, TD = duplication type and D = deletion type. Mb = mega-base-pair. **Single cell whole-genome heatmap:** Genome-wide copy numbers of sequenced aberrant cells. Each row represents a single cell. A total of 96 individual cells were sequenced, and normal cells were excluded from the heatmap. **Representative single cell:** An example whole-genome copy number view of a single cell.

### Figure S2

#### A Case 4 - Parosteal osteosarcoma

circos plot:

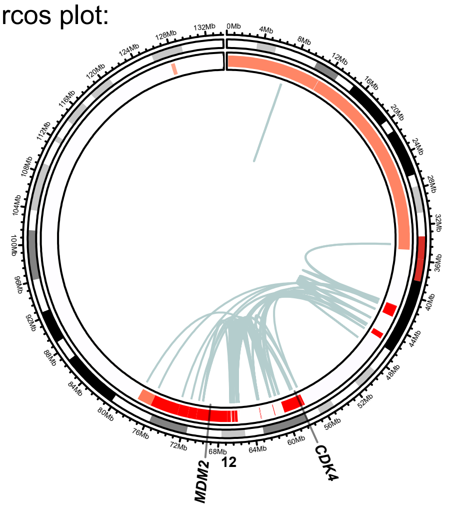


structural variant distribution:

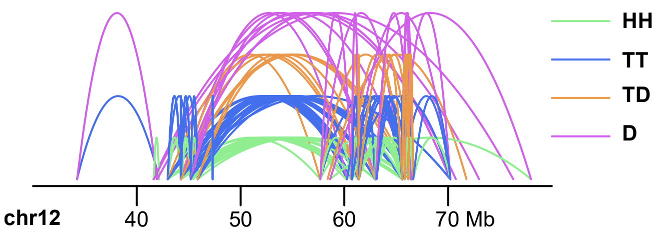


#### B Case 5 - Low-grade central osteosarcoma

Case 5a:
single cell whole-genome heatmap:

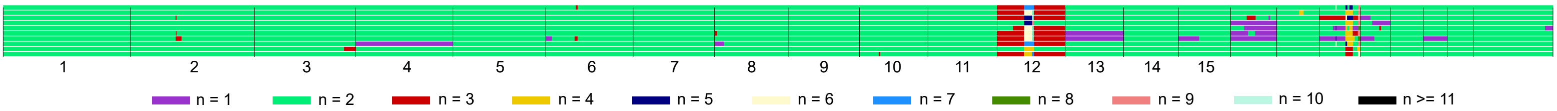


representative single cell:

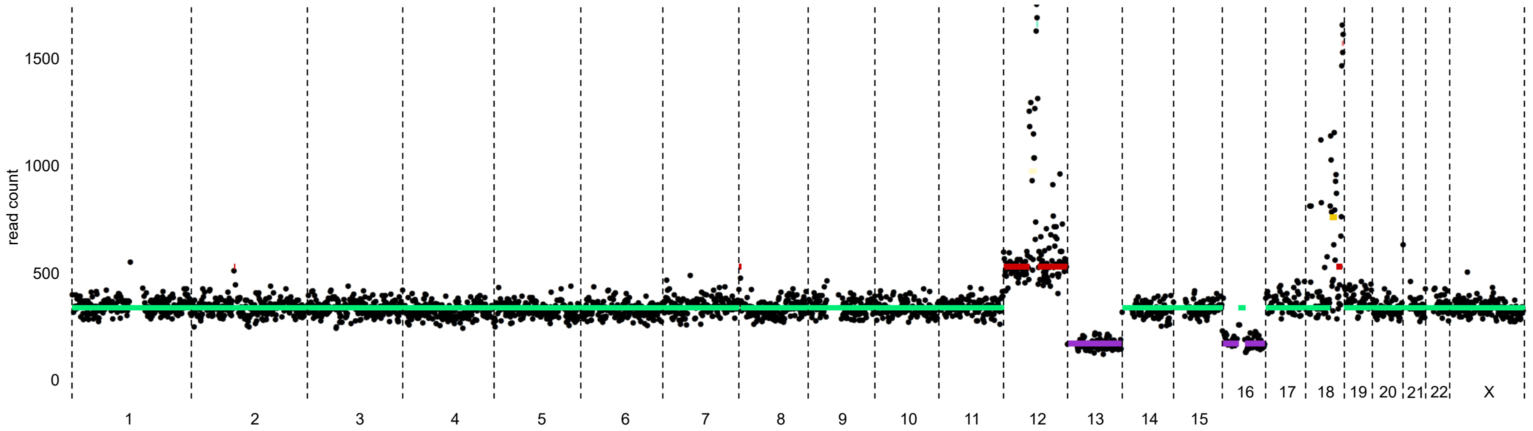


representative single cell:

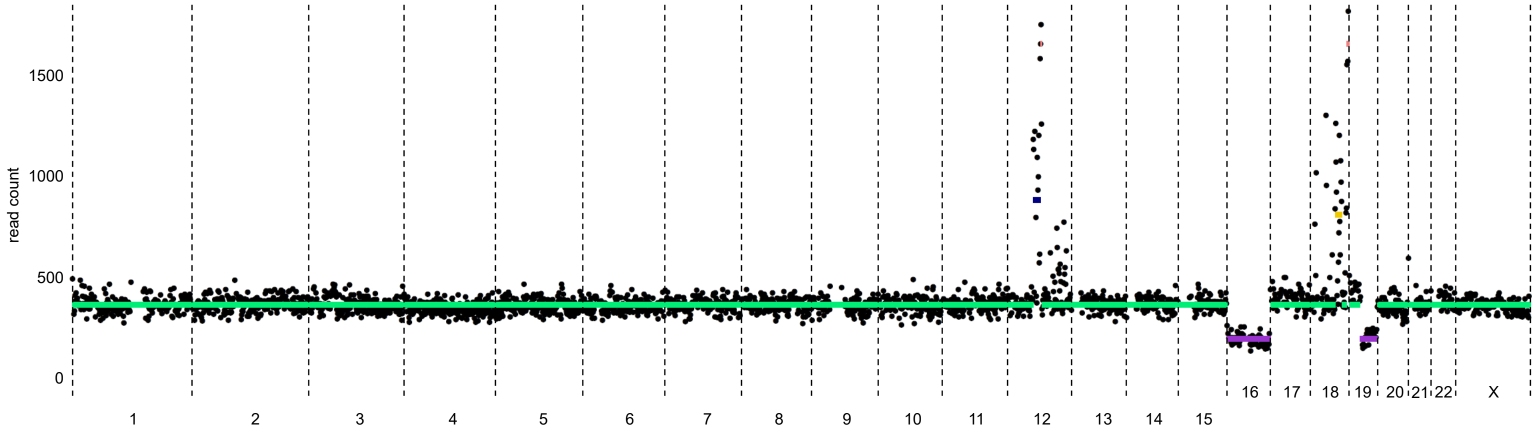


exon coverage plot:

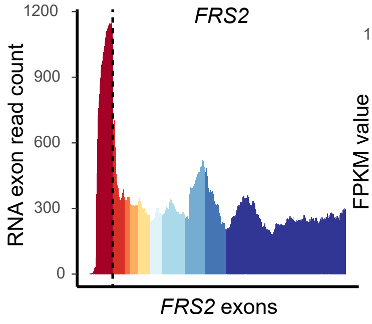


gene expression plot:

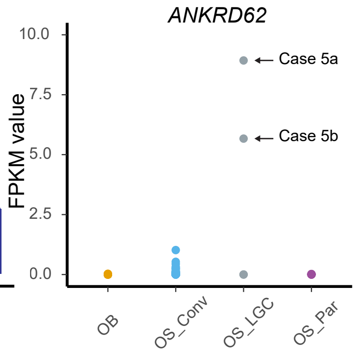


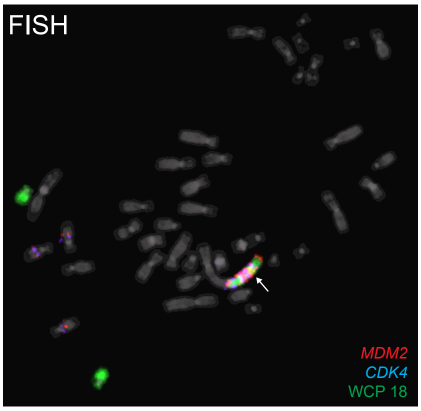

Figure S2
Case 5b:

circos plot:

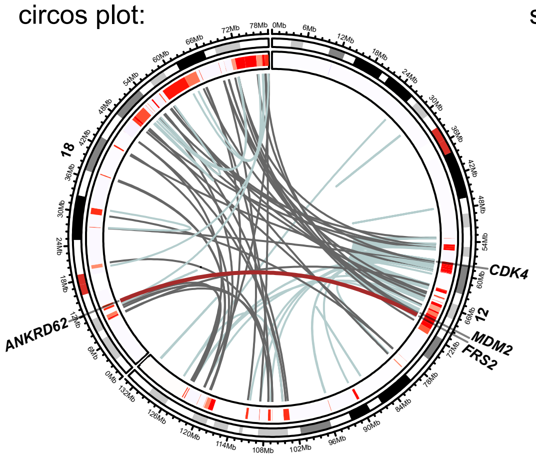


structural variant distribution:

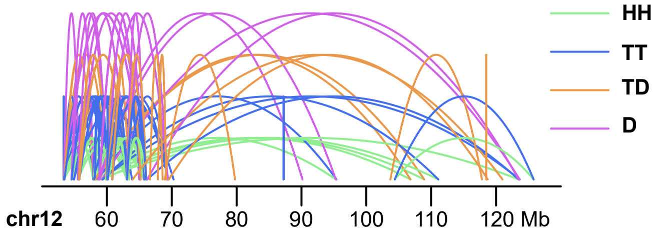


exon coverage plot:

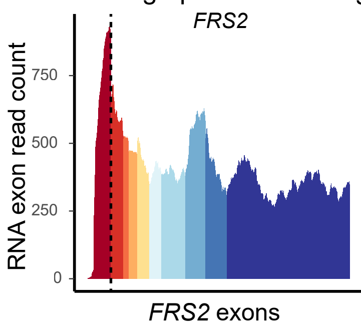


gene expression plot:

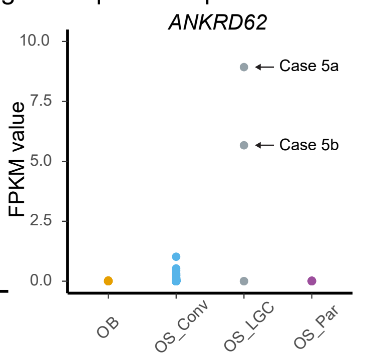


C Case 6 - Parosteal osteosarcoma

circos plot:

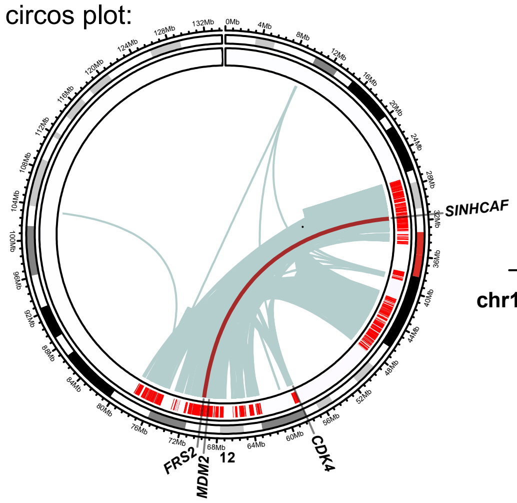


structural variant distribution:

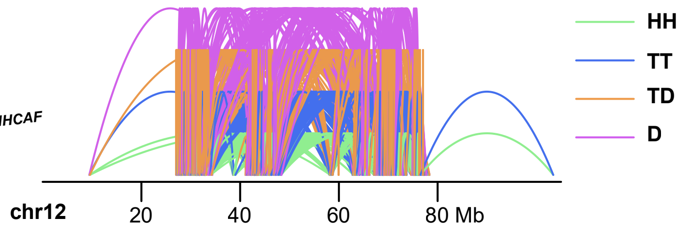


exon coverage plot:

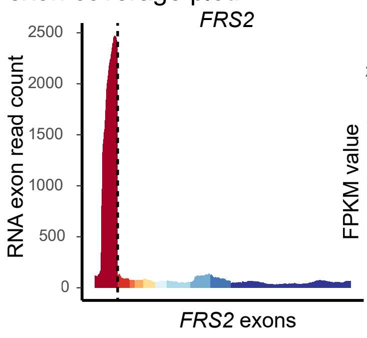


gene expression plot:

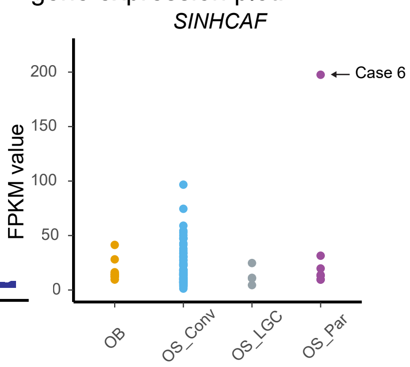


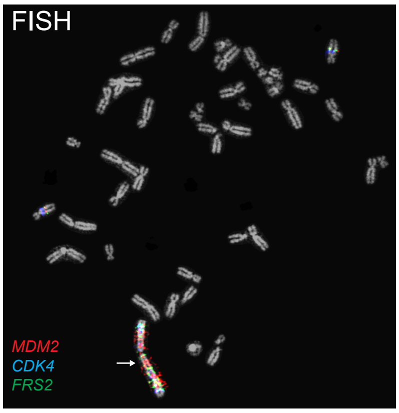

Figure S2

D Case 7 - Dedifferentiated parosteal osteosarcoma

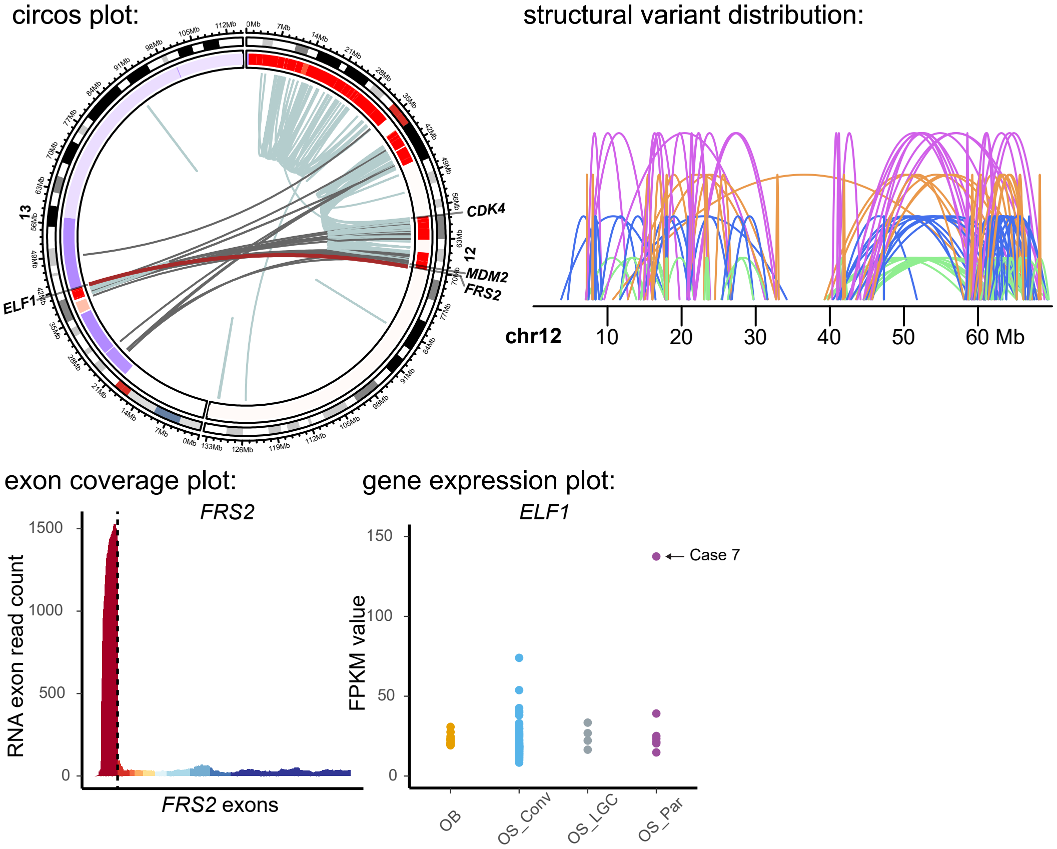


E Case 8 - Parosteal osteosarcoma

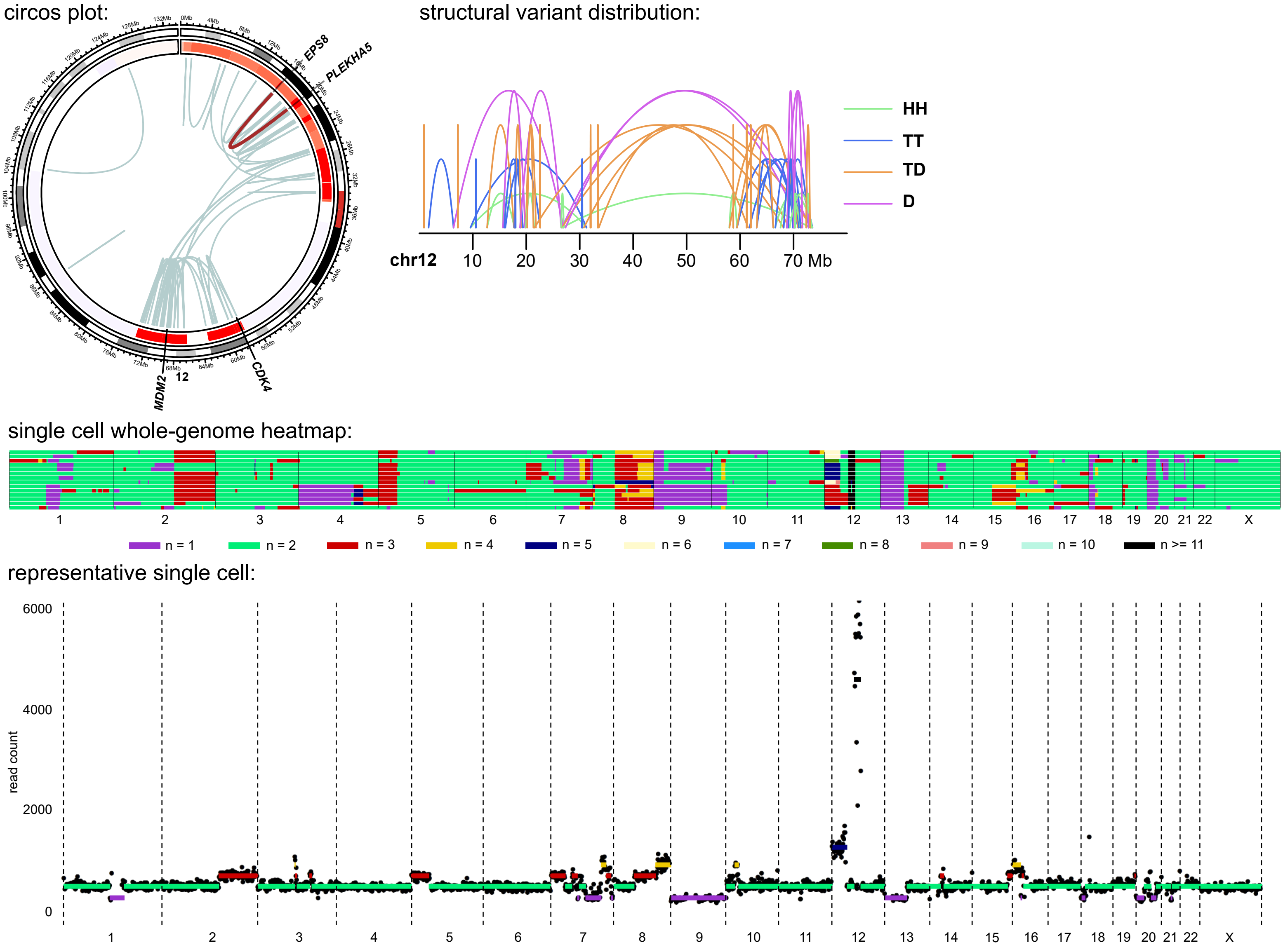

Figure S2

representative single cell:

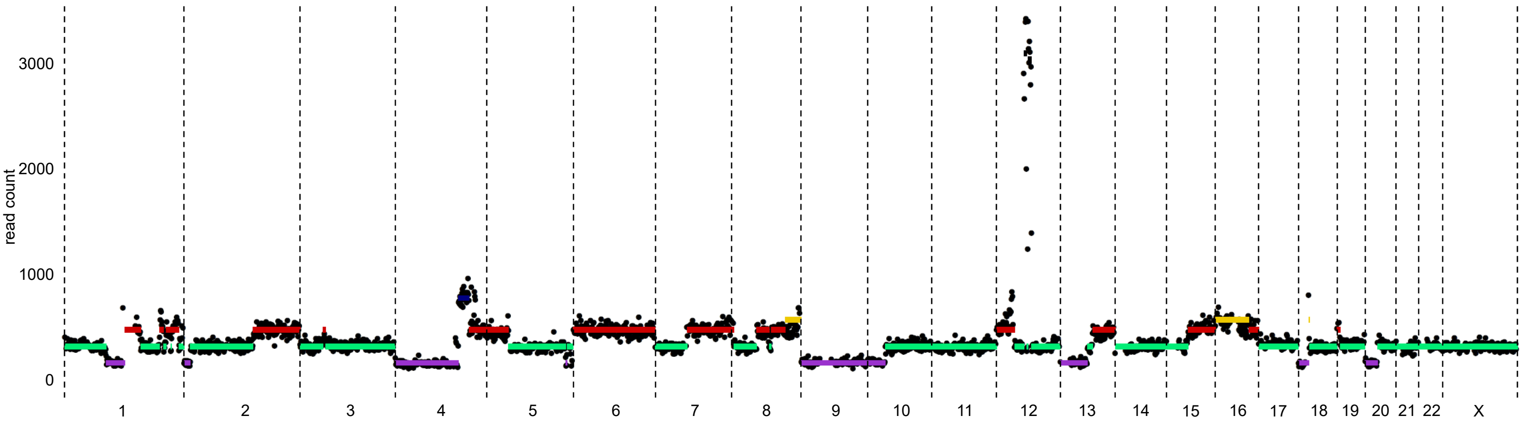


exon coverage plot:

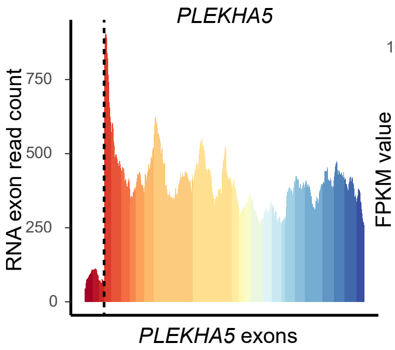


gene expression plot:

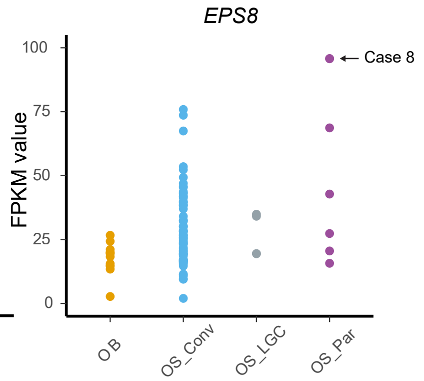


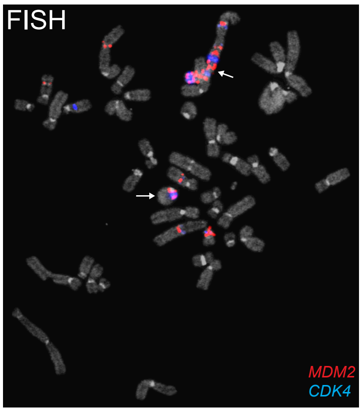


#### F Case 9 - Conventional osteosarcoma

Case 9a:

exon coverage plot:

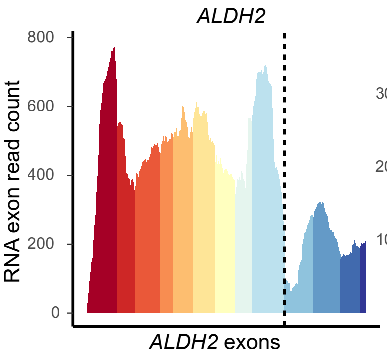


exon coverage plot:

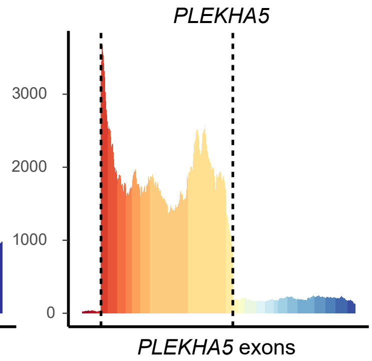


gene expression plot:

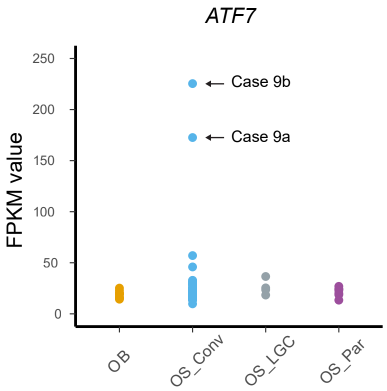


Case 9b:

circos plot:

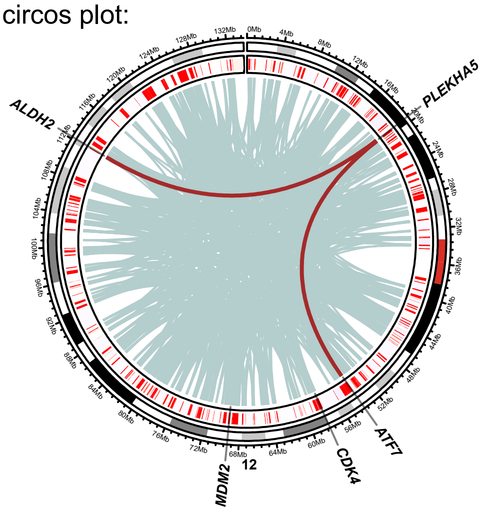


structural variant distribution:

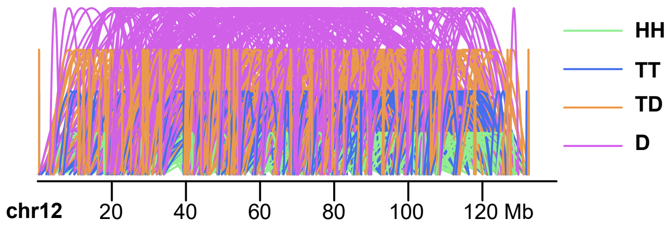

#### Figure S2

single cell whole-genome heatmap:

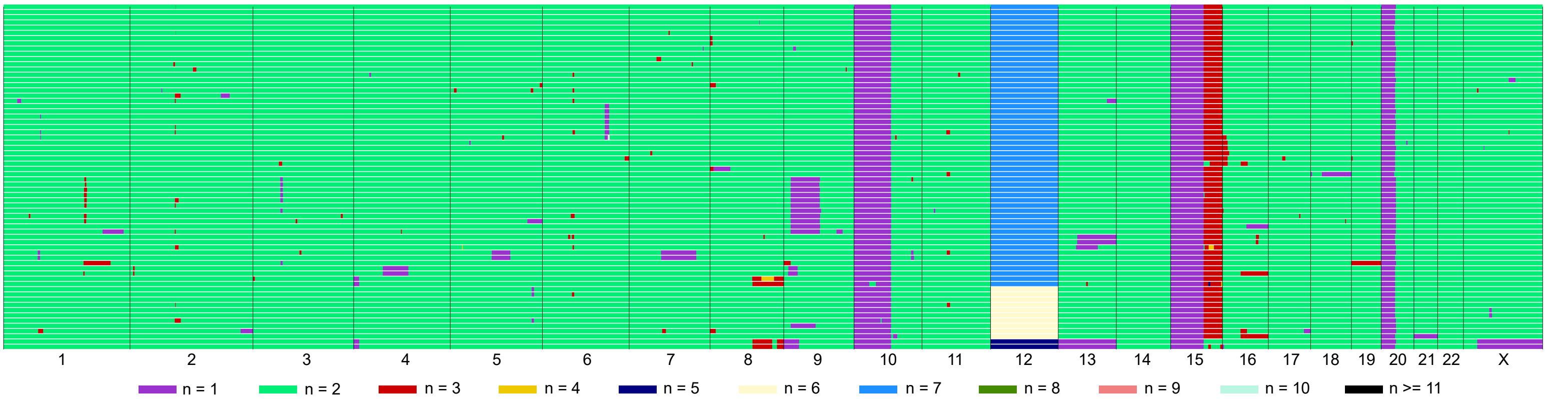


representative single cell:

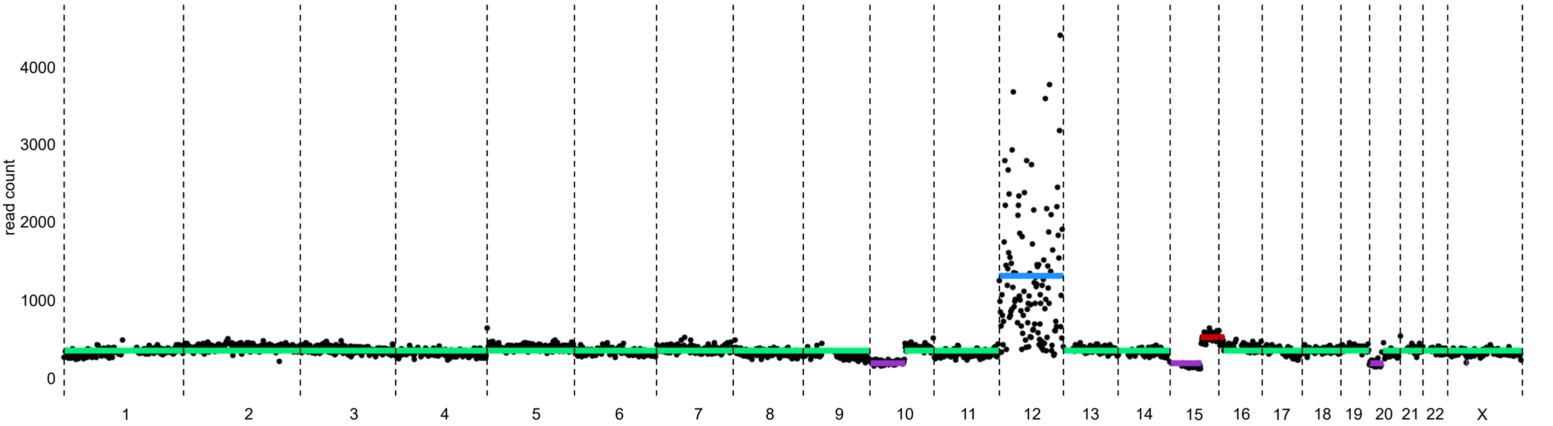


exon coverage plot:

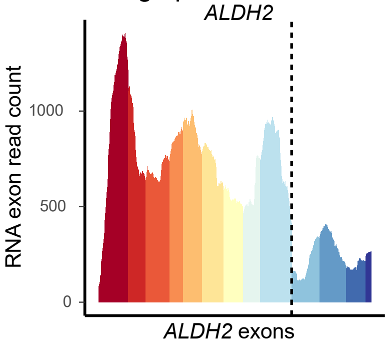


exon coverage plot:

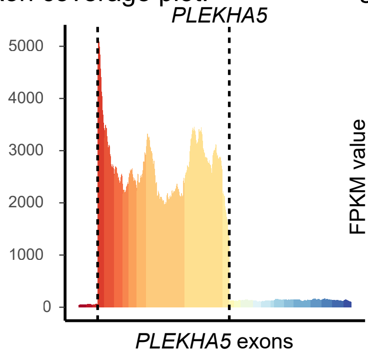


gene expression plot:

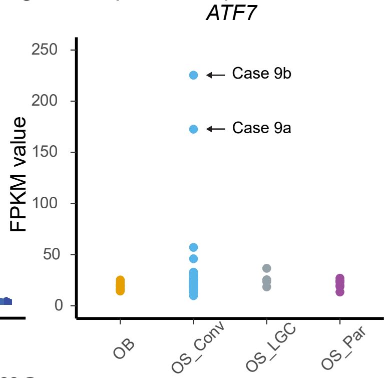


#### G Case 10 - Dedifferentiated parosteal osteosarcoma

circos plot:

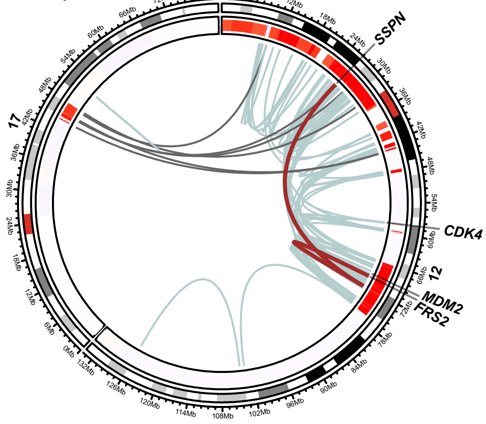


structural variant distribution:

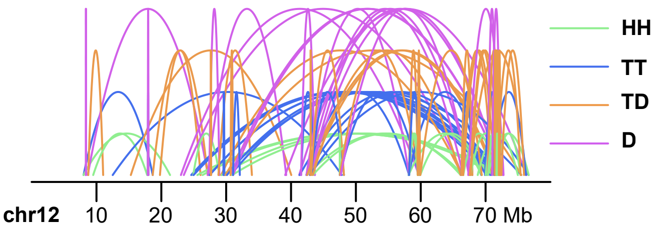


exon coverage plot:

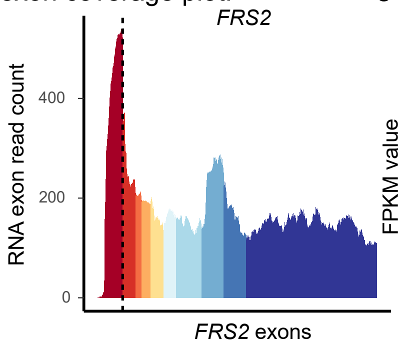


gene expression plot:

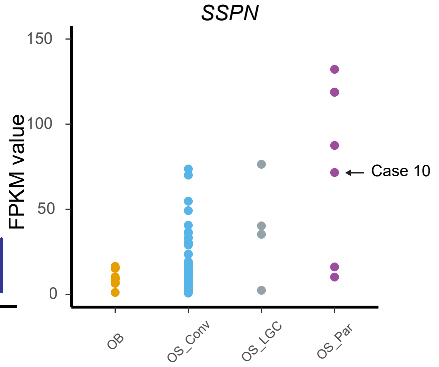

Figure S2

H OS191 - Conventional osteosarcoma

circos plot:

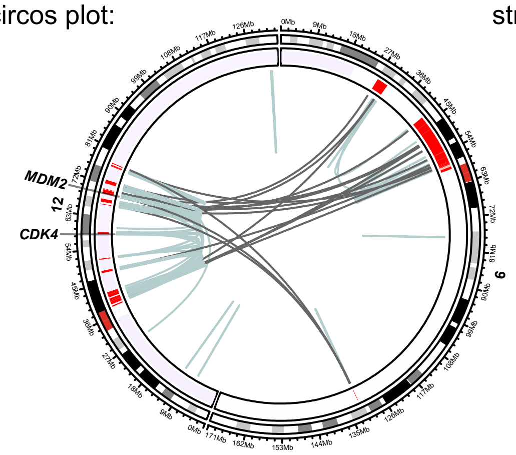


structural variant distribution:

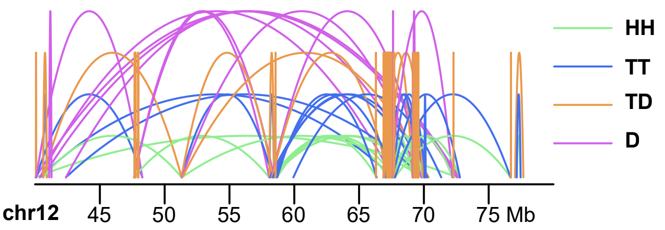


I Case 12 - Dedifferentiated parosteal osteosarcoma

circos plot:

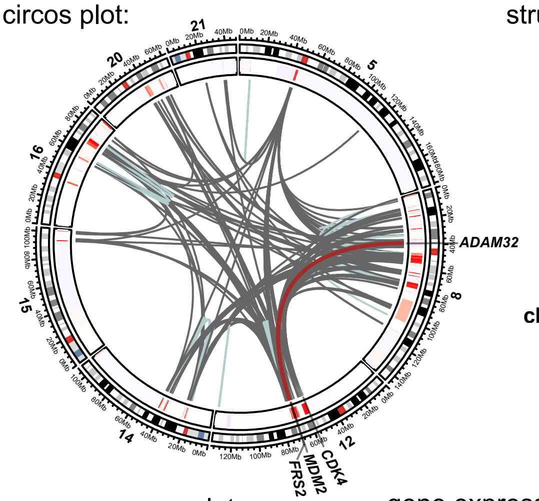


structural variant distribution:

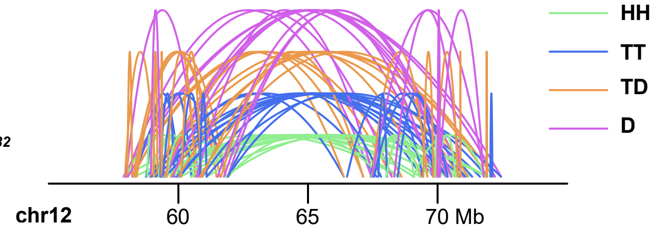


exon coverage plot:

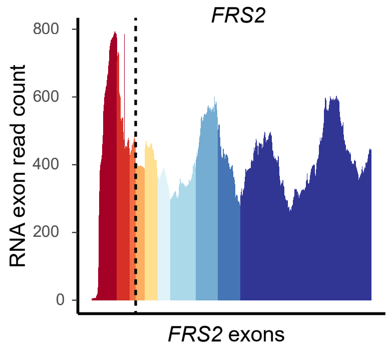


gene expression plot:

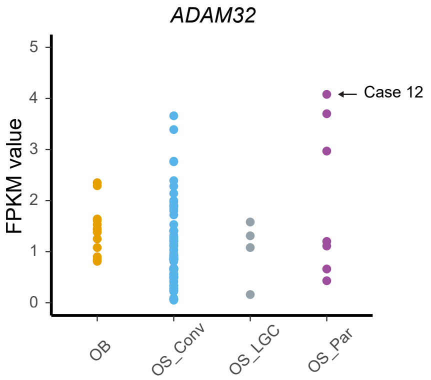


**Figure S2. Group B: combined copy number, structural variant and transcriptomic data. Circos plot:** Red regions in the inner circular track of the circos plot indicate copy number gains, with higher level amplifications being in a more intense shade and lower-level gains in a lighter shade. Blue regions in the inner circular track of the circos plot indicate copy number losses. Intrachromosomal structural variants are depicted in light blue and interchromosomal structural variants in grey. Selected variants are highlighted in brown. **Structural variant distribution:** Intrachromosomal structural variants plotted based on read mapping orientation. Abbreviations: HH = head-to-head inversion, TT = tail-to-tail inversion, TD = duplication type and D = deletion type. Mb = mega-base-pair. **Single cell whole-genome heatmap:** Genome-wide copy numbers of sequenced aberrant cells. Each row represents a single cell. A total of 96 individual cells were sequenced, and normal cells were excluded from the heatmap. **Representative single cell:** An example whole-genome copy number view of a single cell. **Exon coverage plot:** Read coverage per exon of the given gene. Each exon is depicted in a different colour. The dashed line(s) indicates the breakpoint(s) on the RNA level. **Gene expression plot:** Relative gene expression levels of the given gene. The case under study is indicated by an arrow. Abbreviations: OB = osteoblastoma, OS Conv = conventional osteosarcoma, OS LGC = low-grade central osteosarcoma, OS Par = parosteal osteosarcoma (including dedifferentiated parosteal osteosarcoma). **FISH:** Fluorescence in situ hybridization (FISH) was conducted using probes targeting specific regions as indicated in the image.

Figure S3

A Case 13 - Conventional osteosarcoma

circos plot:

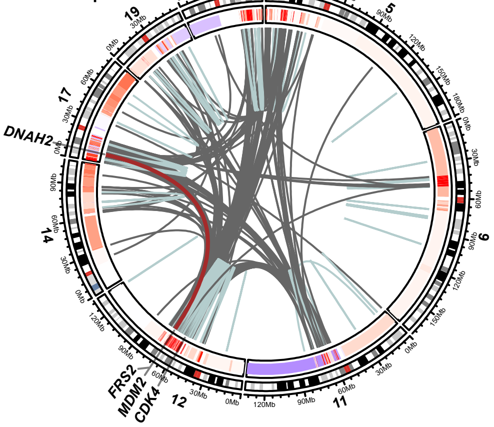


structural variant distribution:

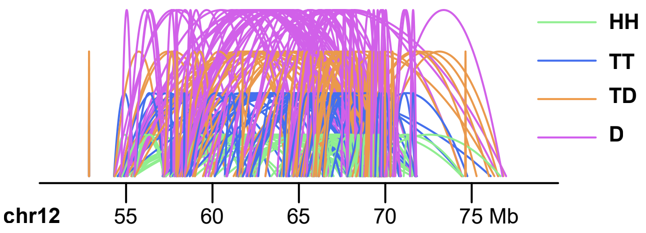


single cell whole-genome heatmap:

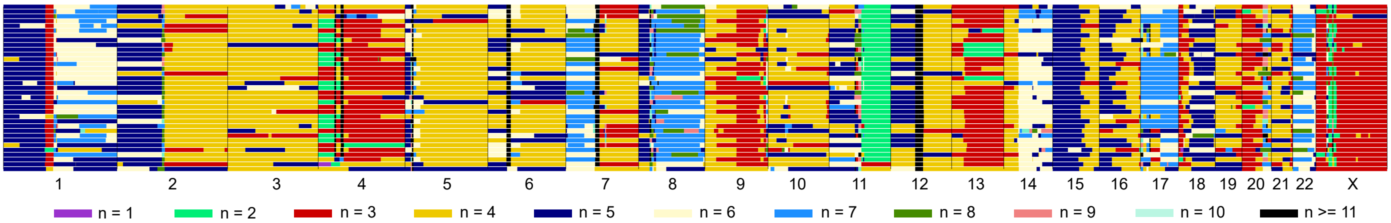


B Case 15 - Conventional osteosarcoma

circos plot:

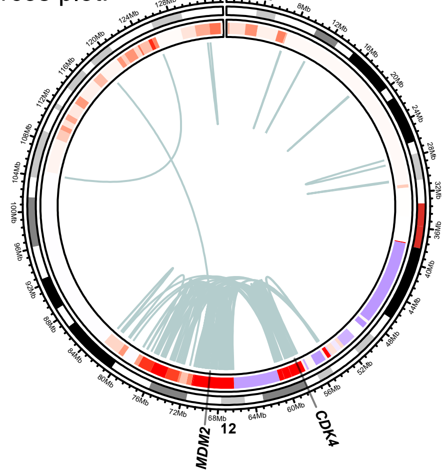


structural variant distribution:

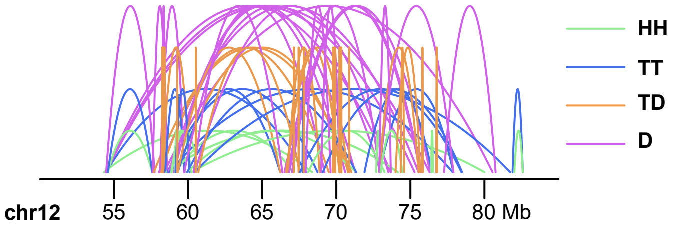


C Case 16 - Conventional osteosarcoma

circos plot:

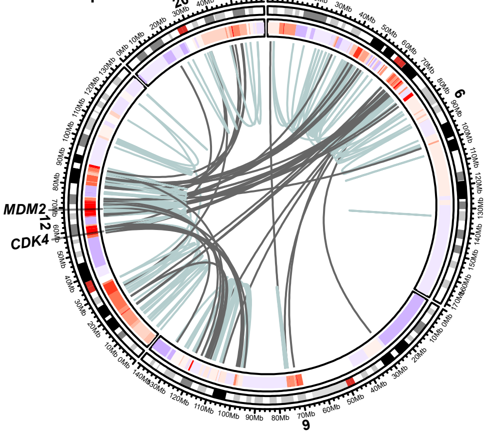


structural variant distribution:

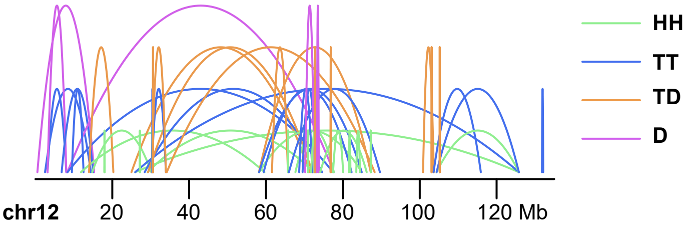

Figure S3

D OS131 - Conventional osteosarcoma

circos plot:

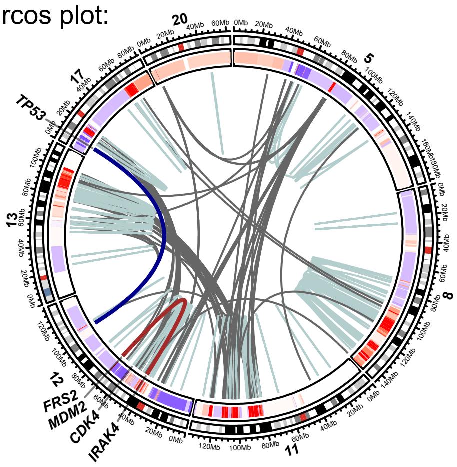


structural variant distribution:

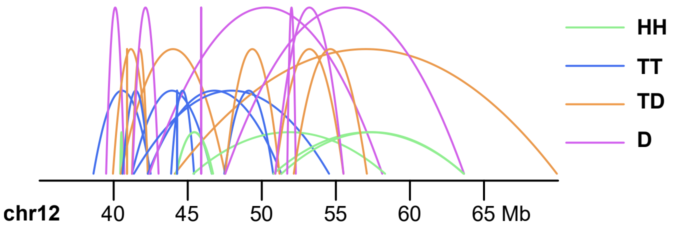


E Case 17 - Conventional osteosarcoma

circos plot:

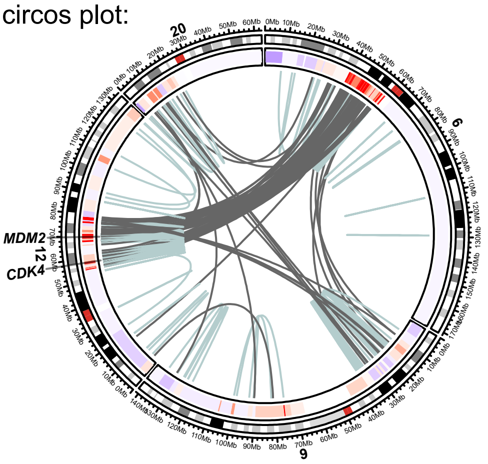


structural variant distribution:

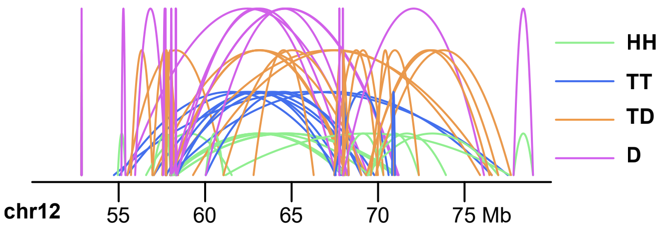


single cell whole-genome heatmap:

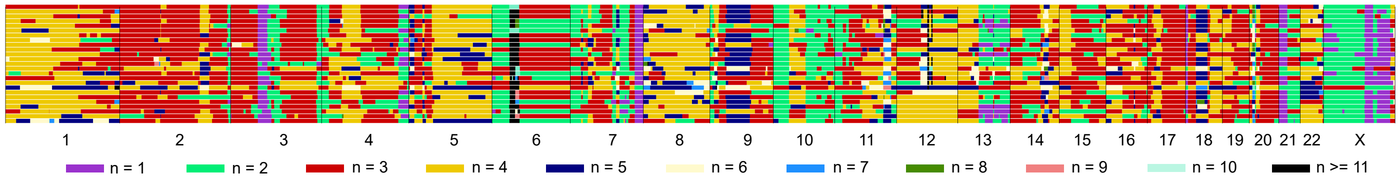


**Figure S3. Group C: combined copy number and structural variant data. Circos plot:** Red regions in the inner circular track of the circos plot indicate copy number gains, with higher level amplifications being in a more intense shade and lower-level gains in a lighter shade. Blue regions in the inner circular track of the circos plot indicate copy number losses. Intrachromosomal structural variants are depicted in light blue and interchromosomal structural variants in grey. Selected variants are highlighted in brown. S**tructural variant distribution:** Intrachromosomal structural variants plotted based on read mapping orientation. Abbreviations: HH = head-to-head inversion, TT = tail-to-tail inversion, TD = duplication type and D = deletion type. Mb = mega-base-pair. **Single cell whole-genome heatmap:** Genome-wide copy numbers of sequenced aberrant cells. Each row represents a single cell. A total of 48 individual cells were sequenced, and normal cells were excluded from the heatmap.

Figure S4

A OS061 - Conventional osteosarcoma

circos plot:

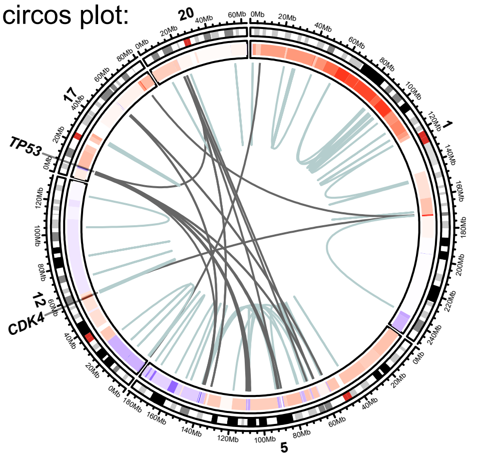


structural variant distribution:

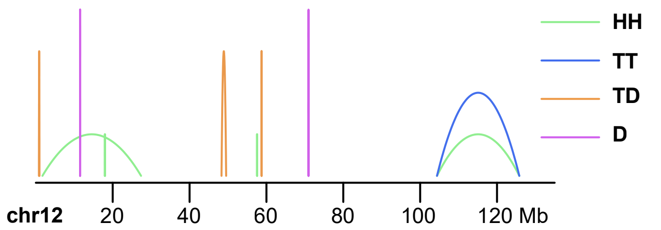


B OS222 - Conventional osteosarcoma

circos plot:

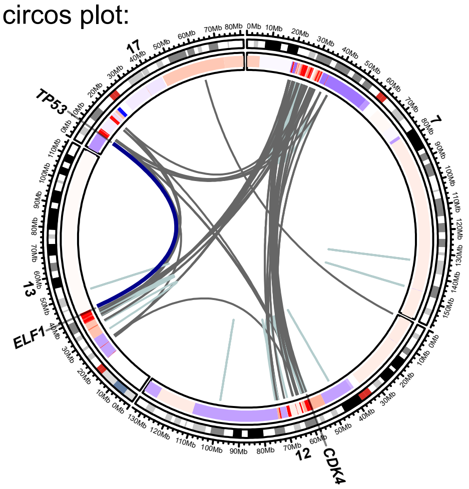


structural variant distribution:

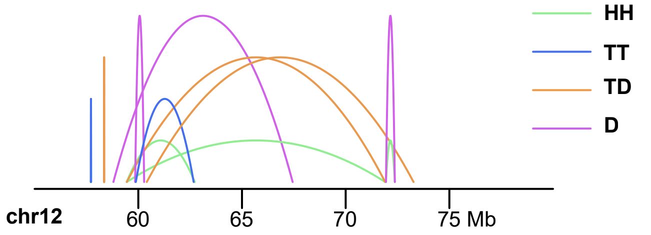


exon coverage plot:

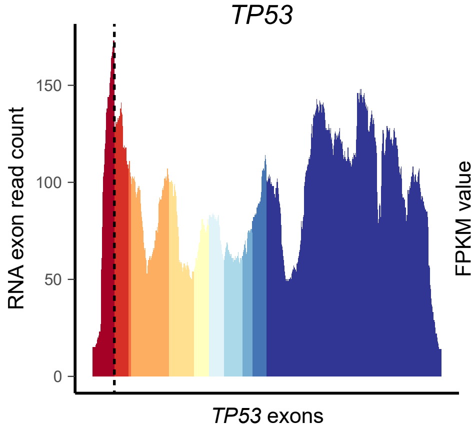


gene expression plot:

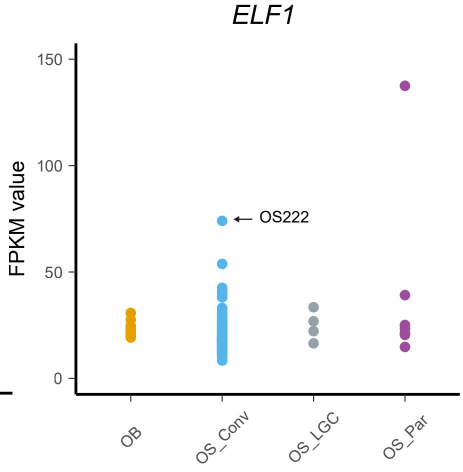


C OS046 - Conventional osteosarcoma

circos plot:

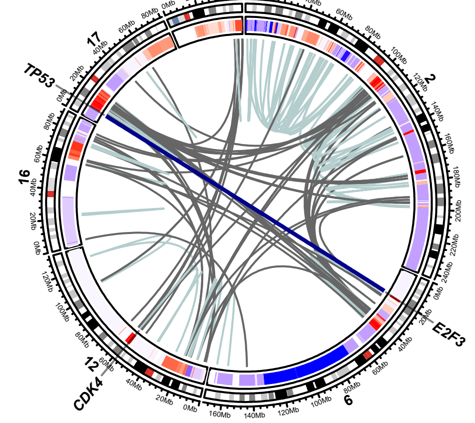


structural variant distribution:

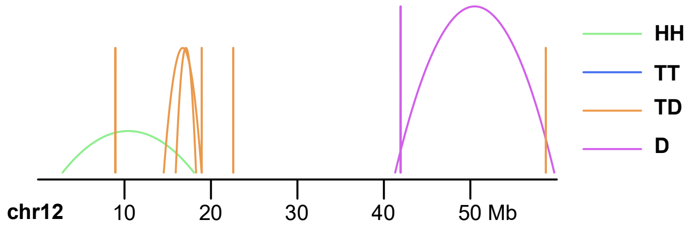

Figure S4

exon coverage plot:

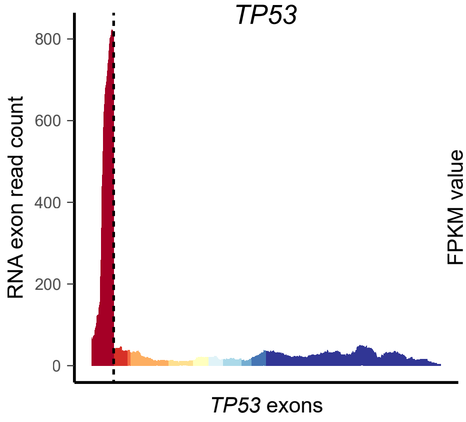


gene expression plot:

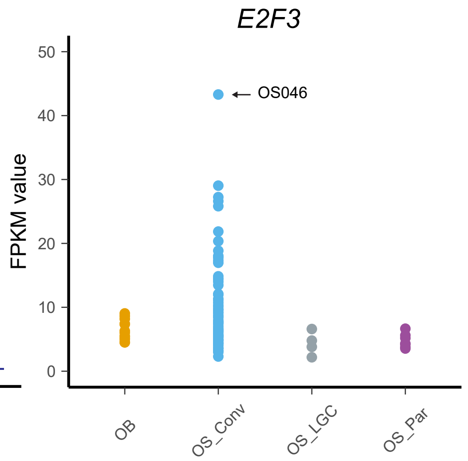


**Figure S4. Group D: combined copy number, structural variant and transcriptomic data. Circos plot:** Red regions in the inner circular track of the circos plot indicate copy number gains, with higher level amplifications being in a more intense shade and lower-level gains in a lighter shade. Blue regions in the inner circular track of the circos plot indicate copy number losses. Intrachromosomal structural variants are depicted in light blue and interchromosomal structural variants in grey. Selected variants are highlighted in dark blue. **Structural variant distribution:** Intrachromosomal structural variants plotted based on read mapping orientation. Abbreviations: HH = head-to-head inversion, TT = tail-to-tail inversion, TD = duplication type and D = deletion type. Mb = mega-base-pair. **Exon coverage plot:** Read coverage per exon of the given gene. Each exon is depicted in a different colour. The dashed line indicates the breakpoint on the RNA level. **Gene expression plot:** Relative gene expression levels of the given gene. The case under study is indicated by an arrow. Abbreviations: OB = osteoblastoma, OS Conv = conventional osteosarcoma, OS LGC = low-grade central osteosarcoma, OS Par = parosteal osteosarcoma (including dedifferentiated parosteal osteosarcoma).

Figure S5

#### Case 1 Group A

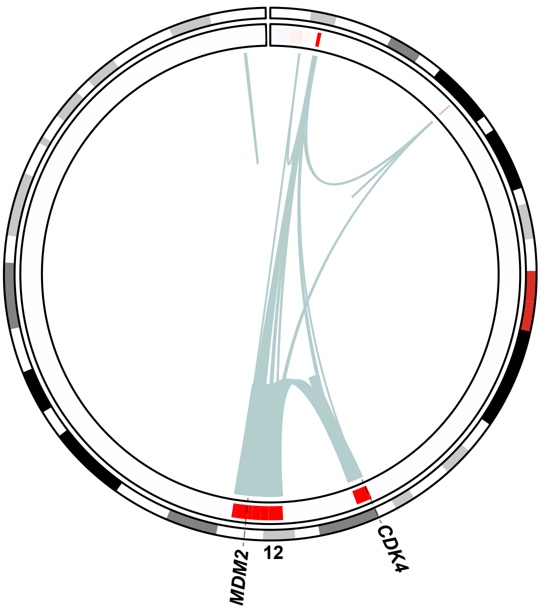


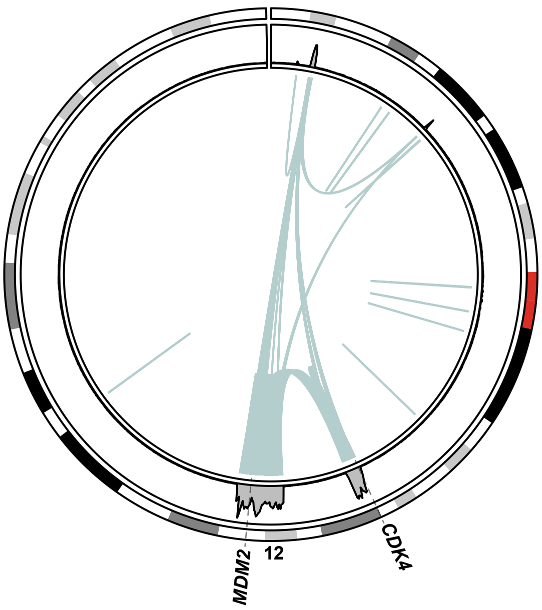


#### Case 2 Group A

Case 2a

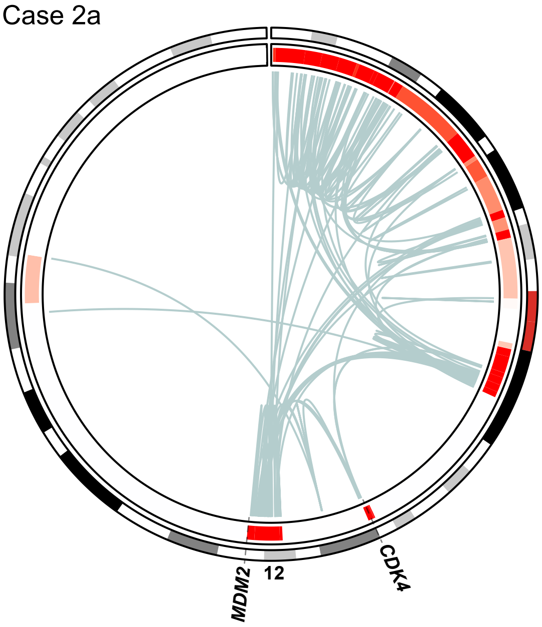


Case 2b

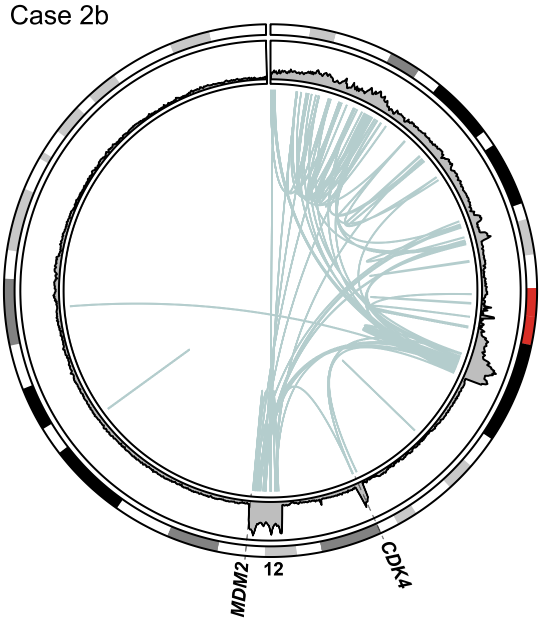


#### Case 7 Group B

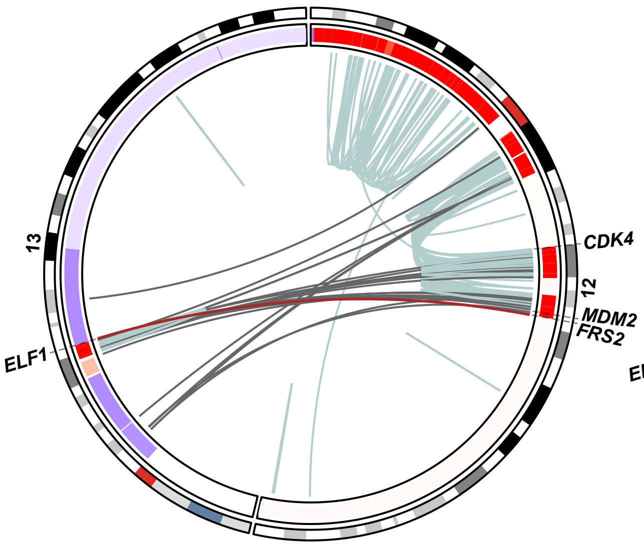


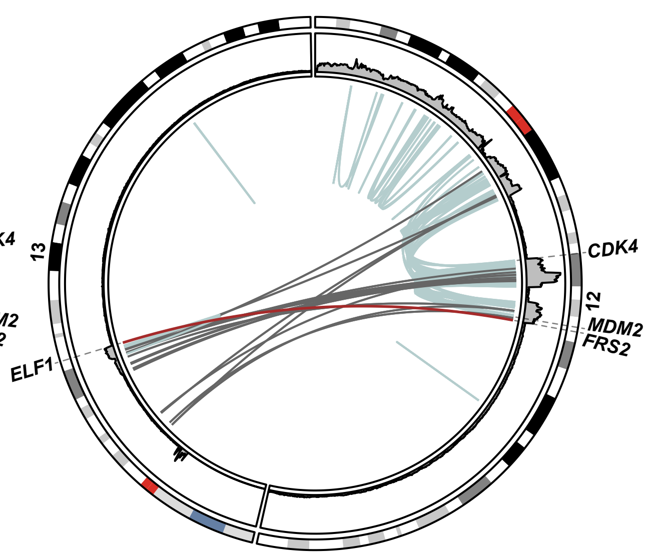

Figure S5

Case 8 Group B

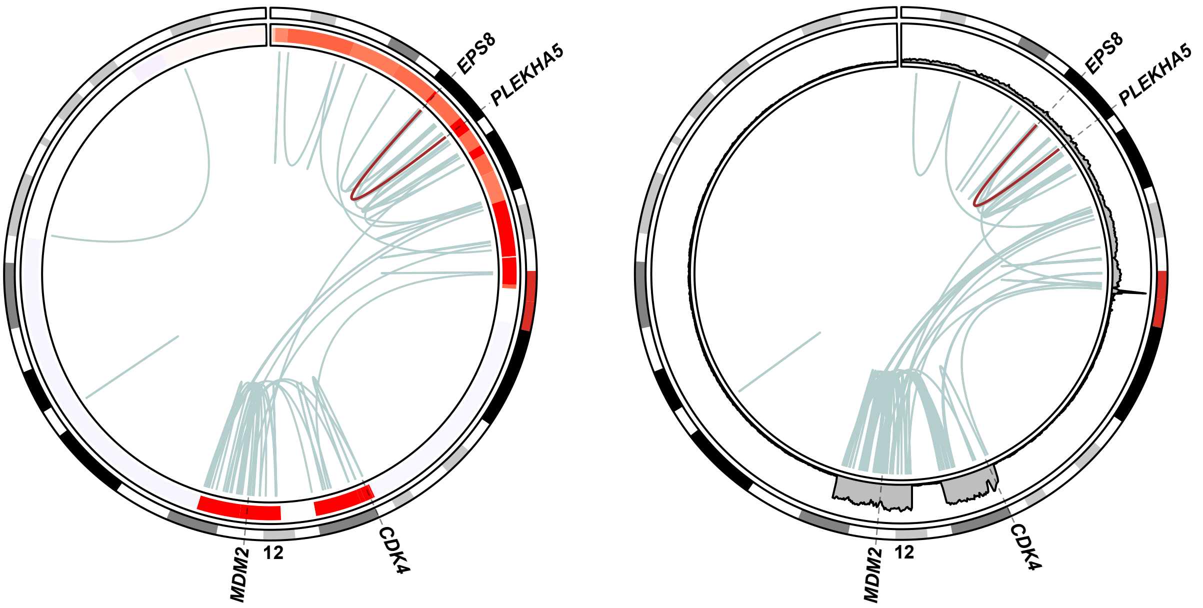


Case 9 Group B

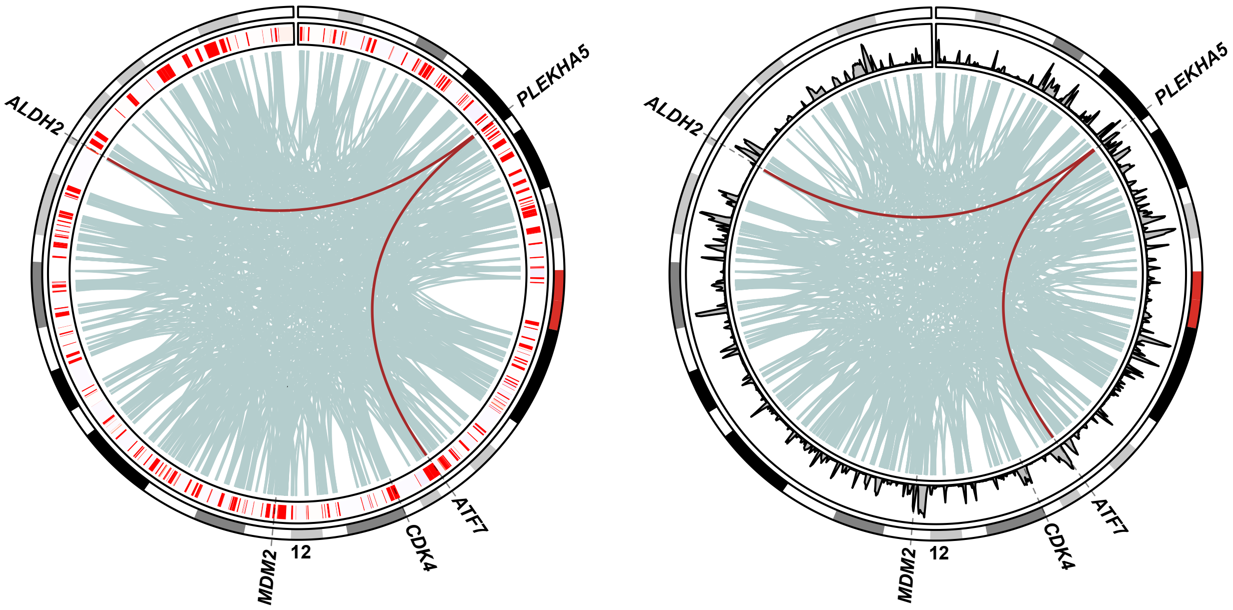


Case 10 Group B

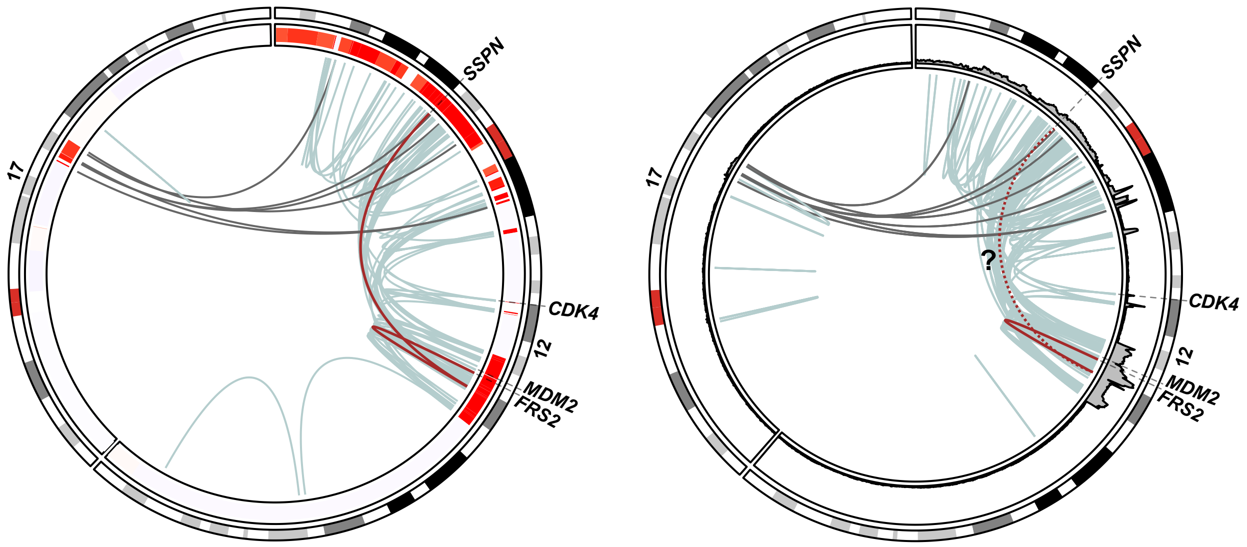

Figure S5

#### Case 12 Group B

Case 12a

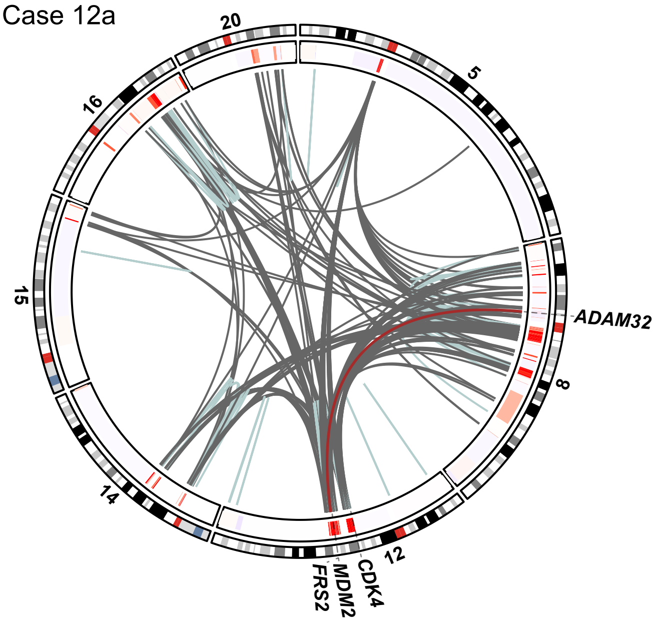


Case 12c

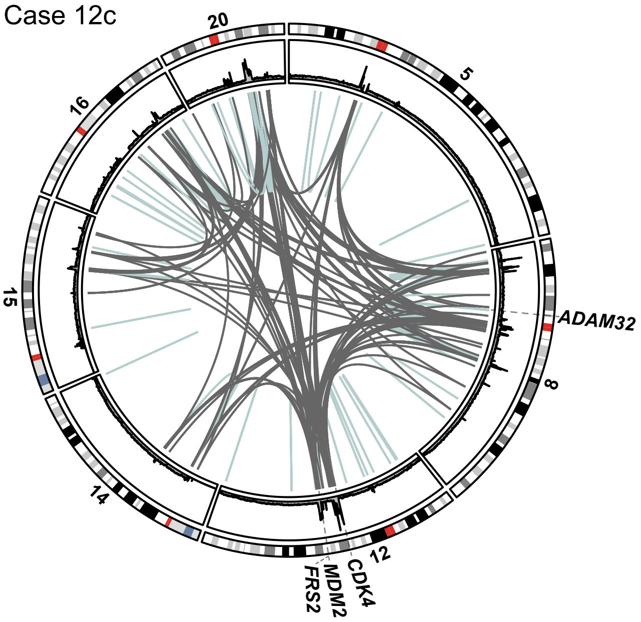


#### Case 13 Group C

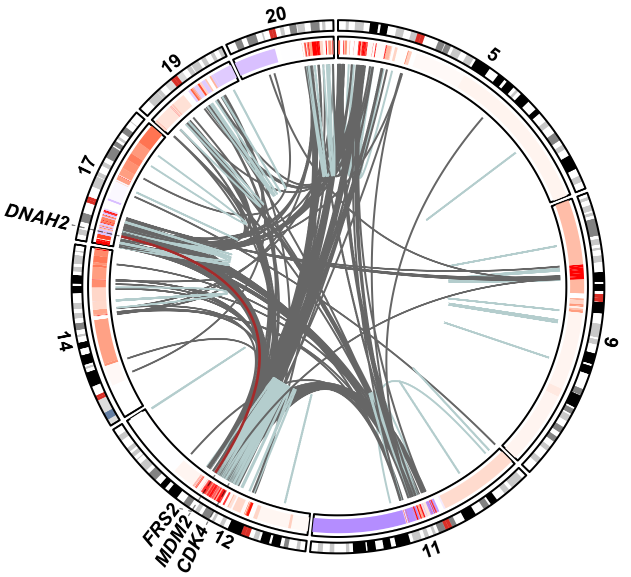


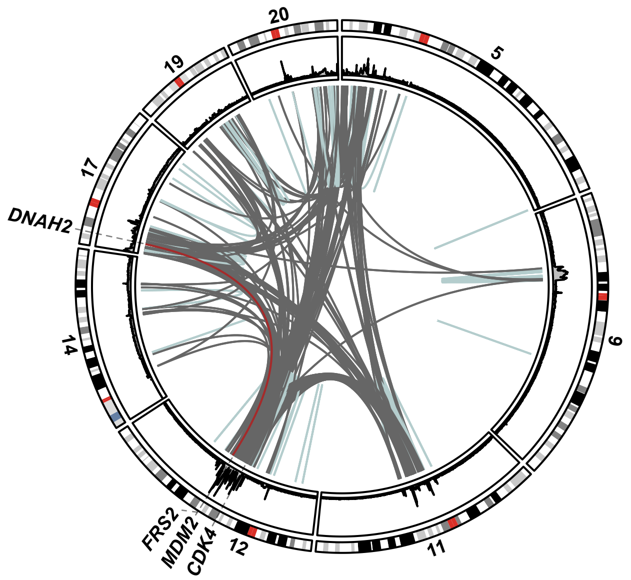


#### Case 14 Group C

Case 14 lacks mate pair sequencing data for comparison.

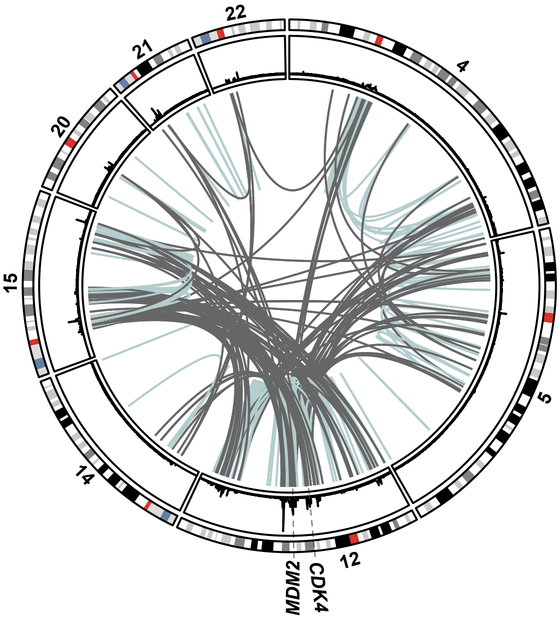

Figure S5

Case 17 Group C

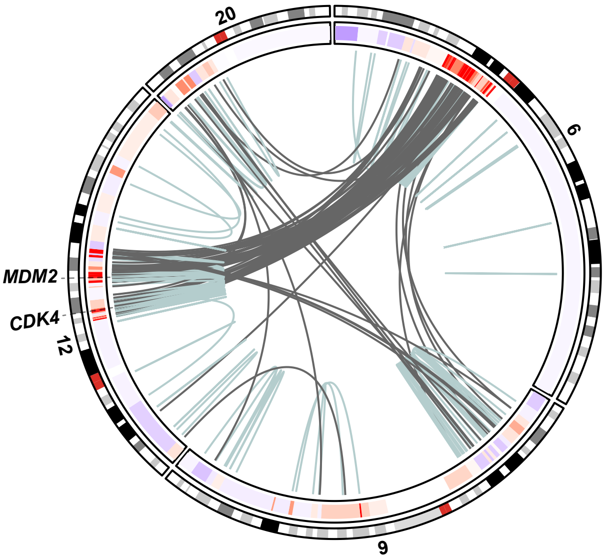


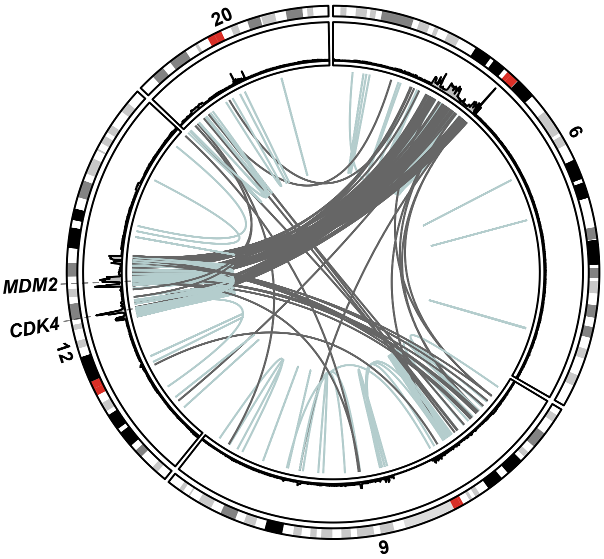


**Figure S5. Side-by-side comparison of mate pair and longread sequencing in selected cases. Left circos plot:** Mate pair sequencing data - Red regions in the inner circular track of the circos plot indicate copy number gains, with higher level amplifications being in a more intense shade and lower-level gains in a lighter shade. Blue regions in the inner circular track of the circos plot indicate copy number losses. Intrachromosomal structural variants are depicted in light blue and interchromosomal structural variants in grey. Selected variants are highlighted in brown. **Right circos plot:** Longread sequencing data – Coverage levels are plotted in the first inner track. Intrachromosomal structural variants are depicted in light blue and interchromosomal structural variants in grey. Selected variants are highlighted in brown.

Figure S6

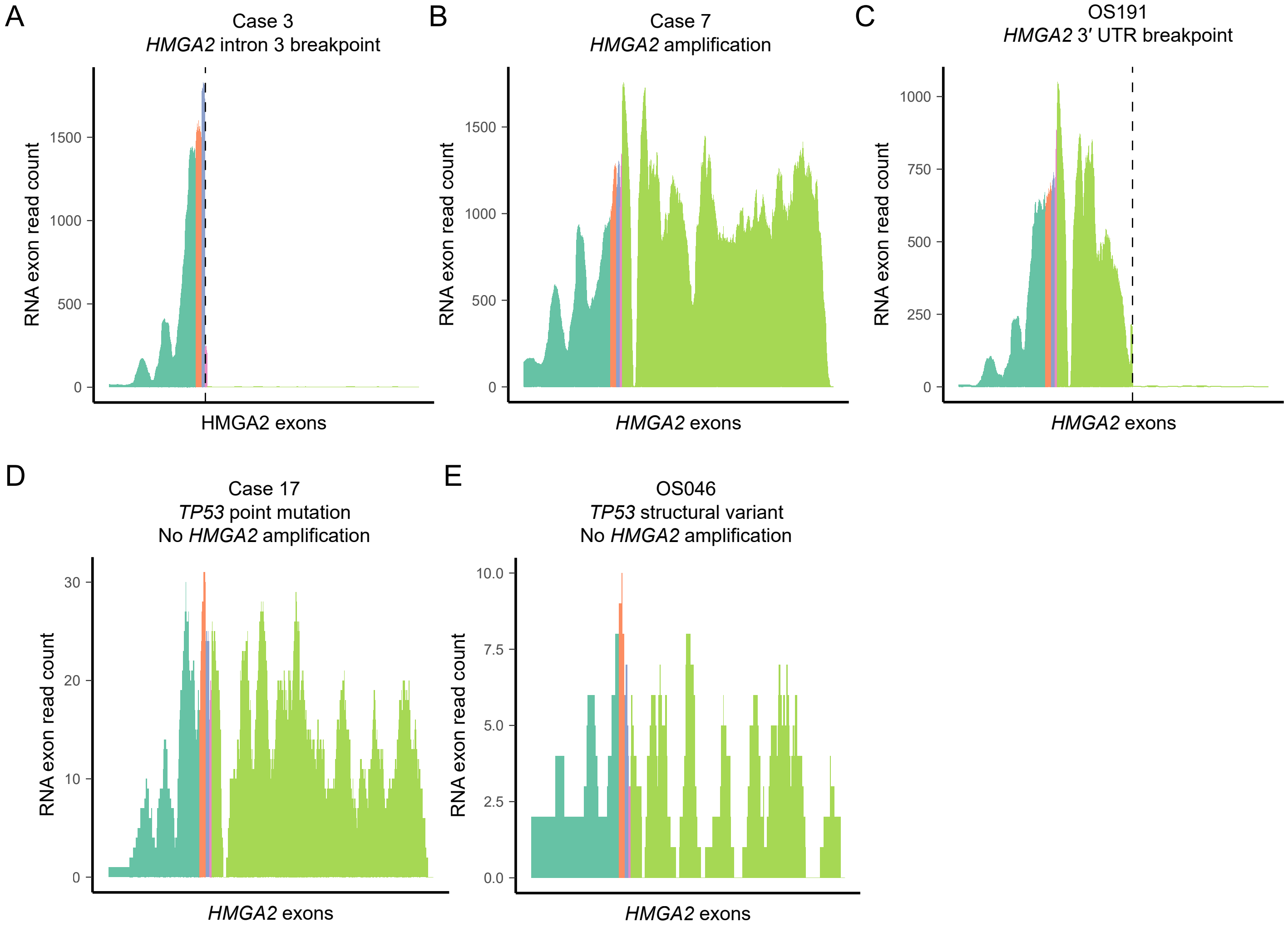


**Figure S6. HMGA2 exon coverage plots in selected cases.** Read coverage per exon of the *HMGA2* gene, with each exon depicted in a different colour. The dashed line indicates the breakpoint on the RNA level if applicable. *HMGA2* is either partially or fully amplified in *TP53*-wildtype cases (Supplementary Table S1) and examples are depicted in **A-C**. Cases with a partial amplification show a sharp decrease in exon coverage 3′ of detected breakpoint. *HMGA2* is not amplified in *TP53*-mutated cases, with examples shown in **D** and **E**.

Figure S7

A B

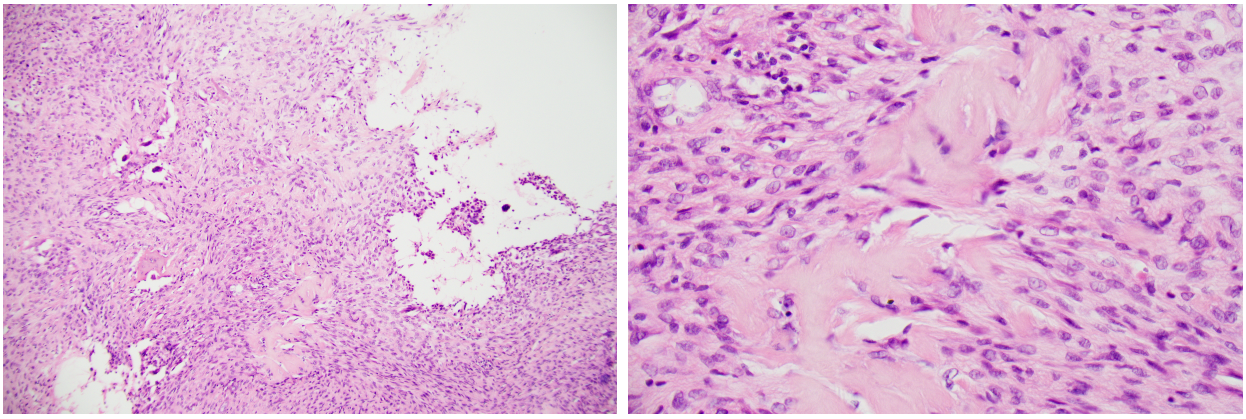

**Figure S7. Histological re-evaluation of Case 1.** Photomicrographs of haematoxylin-eosin stained decalcified tumour tissue at x100 **(A)** and x400 **(B)** times magnification. The tumour consisted of moderately cellular fascicules of spindle cells with mild atypical features. Mitotic figures were rare. Throughout the tumour, neoplastic irregular bone could be identified **(B)**. The morphology was consistent with a low-grade central osteosarcoma.
